# Genomic prediction of resistance to *Phytophthora megakarya* in cacao from a sparse host-pathogen screen

**DOI:** 10.64898/2026.11.03.764016

**Authors:** Ina Schlathoelter, Andrew J. Gitto, Adriana L. Arciniegas, Mariana Herrera-Corzo, Oluwaseun Olawale Kolawole, Erica M. Goss, Donald Livingstone, Alana Firl, Donna R. Henderson, Derek R. Drost, Dayana Rodezno, Jean-Philippe Marelli, Jeremy T. Brawner

## Abstract

Black pod rot caused by *Phytophthora megakarya* is a major disease impacting cacao (*Theobroma cacao* L.) production in West Africa. Resistance is influenced by genetic variation in both host and pathogen populations. While genomic prediction approaches have been widely applied in plant breeding, most studies incorporate only host genomic information. The value of explicitly modeling pathogen genomic relatedness in quantitative disease prediction remains largely unknown.

We genotyped 947 clones from the International Cacao Collection at CATIE using a 10K-SNP panel and characterized 67 *P. megakarya* isolates by sequencing 3,505 virulence-associated genes and 2,774 SNP loci. Cacao clones were extensively admixed, while isolates formed three geographically structured subpopulations. Mixed linear models were fitted to lesion diameters on detached pods of 350 genotyped clones inoculated with 58 isolates in 30 bioassays, covering 5% of possible clone-by-isolate combinations.

Differences among isolates explained a larger share of lesion-size variation than differences among clones (0.34 and 0.27 of phenotypic variance, respectively). After accounting for broad pathogen subpopulation structure, incorporating within-subpopulation genomic relatedness among isolates improved neither model fit nor prediction, whereas incorporating genomic relationships among cacao clones improved model fit and enabled prediction of phenotypically untested clones with a predictive ability of 0.41. Clone rankings were stable across pathogen subpopulations (genomic correlation 0.86 ± 0.06), whereas clone-by-isolate interactions accounted for little phenotypic variation and were not resolved with confidence in the sparse design. An exploratory ancestry analysis associated several Upper Amazon ancestry components with smaller predicted lesions.

These are the first genomic predictions of resistance to *P. megakarya* in cacao. In this strongly structured host-pathogen system, genomic relationships among hosts carried most of the predictable signal beyond broad pathogen subpopulation effects, and the predictions provide a starting point for prioritizing screening and selection efforts.

## Introduction

Cacao (*Theobroma cacao* L.) beans are the raw material of the global chocolate industry, valued at approximately US$167 billion in 2024 (IMARC Group, 2024). Originating in the upper Amazon region of South America (Zarrillo et al., 2018), cacao subsequently became established as a major crop throughout tropical regions worldwide. Africa accounts for approximately 70% of global cocoa production, with Côte d’Ivoire and Ghana together contributing nearly half of the global supply (International Cocoa Organization, 2025). Cocoa production is predominantly carried out by smallholder farmers, with more than two million producers estimated in Africa (Kissi & Herzig, 2024). Cacao is a key cash crop in the humid tropics, supporting the livelihoods of millions of people worldwide, historically estimated at 40–50 million (CacaoNet, 2012).

Cacao production is threatened by several diseases. Black pod rot, caused by several species of the genus *Phytophthora*, is one of the most destructive biotic threats to cacao and accounts for half of reported global yield losses (Marelli et al., 2019). Among the causal agents, *Phytophthora megakarya* is particularly aggressive and currently restricted to West and Central Africa (Wessel & Quist-Wessel, 2015). It can result in up to 90% yield loss without control (Bailey et al., 2016; Nyassé et al., 1999). The oomycete pathogen most likely originated in Cameroon (Mfegue et al., 2012; Nyassé et al., 1999) and subsequently spread westward through West Africa, reaching Côte d’Ivoire by the early 2000s (Akrofi et al., 2015). While *P. megakarya* has largely displaced *P. palmivora* as the dominant black pod pathogen in Cameroon and Nigeria, both species continue to occur in Ghana and Côte d’Ivoire (Ali et al., 2016, 2017). Potential biological advantages contributing to the competitive success of *P. megakarya* include earlier and more abundant sporulation, rapid appressorium formation facilitating infection of unwounded pods, and prolonged survival in soil (Ali et al., 2016; Bailey et al., 2016; Ward & Griffin, 1981). The continued expansion of *P. megakarya* across West and Central Africa presents significant challenges for sustainable disease management.

An important effort to make cacao production more sustainable and resilient to diseases is the development of resistant varieties through breeding. As in other perennial crops, genetic improvement in cacao is slow. With a juvenility phase of 3-5 years, a single selection cycle can take more than a decade. Two or more cycles are usually necessary before releasing a new variety (Kulesza et al., 2024). Breeding for durable disease resistance is further complicated by genetically structured pathogen populations, in which isolates can differ in aggressiveness but are rarely characterized alongside the host material. Understanding the genetic determinants underlying host resistance and pathogen virulence is therefore essential.

Resistance to *Phytophthora* spp. in cacao is predominantly quantitative and polygenic, with a substantial additive genetic component (Ali et al., 2017; Nyassé et al., 2007). Although cacao genotypes differ considerably in their susceptibility to *P. megakarya*, complete immunity has not been demonstrated, and resistance is generally expressed as reduced disease severity (Nyassé et al., 2007; Pokou et al., 2019). Large *ex situ* collections, such as the International Cacao Collection (IC3) at the Centro Agronómico Tropical de Investigación y Enseñanza (CATIE), conserve diverse cacao genetic resources and provide an important platform for identifying and characterizing novel sources of resistance (Engels, 1981; Fister et al., 2020; Mata-Quirós et al., 2018). These collections encompass representatives of the major genetic groups of cacao (Motamayor et al., 2008; Todd et al., 2025; Zhang et al., 2012) and capture both distinct ancestral lineages and extensive admixture. Comprehensive disease phenotyping of large germplasm collections is logistically challenging and expensive, particularly when the pathogen is a quarantine organism that must be handled under containment away from the collection site. In those cases, only a limited subset of host–isolate combinations can realistically be assayed. Being able to confidently use those sparse, unbalanced datasets would substantially accelerate the discovery of potential new sources of disease resistance and guide targeted screening efforts.

Genomic prediction comprises a family of methods that rely on the association between phenotypes and genotypes in a training population to predict genotype performance in a related set of individuals from genetic markers alone (Meuwissen et al., 2001). It does not rely on the direct identification of causal polymorphisms and can be highly effective for selecting on polygenic traits or traits with significant genotype-by-environment interaction. In plant breeding, genomic prediction approaches have demonstrated considerable potential for predicting complex quantitative traits, including disease resistance, and may improve breeding efficiency by enabling the selection of genotyped individuals without direct phenotypic evaluation (Enciso-Rodriguez et al., 2018; Lorenz et al., 2012; Poland & Rutkoski, 2016; Rolling et al., 2020; Rutkoski et al., 2014).

Although genomic prediction is widely used in plant and animal breeding programs, its application in cacao remains limited (Baek et al., 2025, 2026; Bekele et al., 2022; McElroy et al., 2018; Romero Navarro et al., 2017). A genomic-prediction study of black pod resistance in cacao evaluated resistance under low natural *P. palmivora* disease pressure (Romero Navarro et al., 2017). They evaluated 148 clones for eight years under natural infection and estimated a broad-sense heritability of 0.52 and genomic prediction accuracy ranging from 0.46 to 0.51 (Romero Navarro et al., 2017). A second study evaluated genomic prediction of black pod rot resistance in a population of 495 cacao progeny derived from eight interconnected biparental crosses at CATIE, using artificial pod inoculations with *P. palmivora* (Jacquotte et al., 2026). Genomic prediction achieved predictive abilities of 0.45–0.46 under fivefold cross-validation, which decreased to 0.36 when entire families were excluded from the training population. The study further demonstrated that predictive ability increased with training population size, whereas gains from additional SNP markers were minimal beyond approximately 1,000 markers (Jacquotte et al., 2026).

Furthermore, the explicit incorporation of genomic information from both the host and the pathogen remains uncommon in plant disease prediction (Domínguez-Rondón et al., 2025; Hudson et al., 2024; Hudson & Brawner, 2025). Hudson et al. (2024) integrated the pathogen genetic relationship matrix through reaction norm models (Jarquín et al., 2014) and dual-genome similarity matrices, which provide a means to partition phenotypic variance into components attributable to host genetic effects, pathogen genetic effects, and their interaction. This strategy was applied to *Fusarium* ear rot in maize and *Septoria tritici* blotch in wheat. In both systems, pathogen genotypes explained part of the variation (Domínguez-Rondón et al., 2025; Hudson et al., 2024; Hudson & Brawner, 2025). Because pathogen genetic variation can substantially contribute to disease phenotypes, jointly modeling genomic relationships in both the host and the pathogen is an appealing extension of conventional host-only genomic prediction.

Recent genomic resources for *P. megakarya*, including a reference genome assembly and annotation of infection-related gene families such as secreted RxLR effectors, Crinkler proteins, necrosis-inducing proteins, proteases, detoxification enzymes, and carbohydrate-active enzymes, provide new opportunities to characterize pathogen diversity at high resolution (Morales-Cruz et al., 2020). Population-level genomic analyses have revealed pronounced genetic structure within *P. megakarya*, often associated with geographic origin (Mfegue et al., 2012; Morales-Cruz et al., 2020; Nyassé et al., 1999). However, the extent to which genomic differentiation among pathogen subpopulations translates into variation in virulence, and thereby influences quantitative host resistance, remains uncharacterized.

Here, we integrate genome-wide SNP data from the IC3 cacao collection with target capture-based sequencing of virulence-associated genomic regions in a geographically diverse panel of *P. megakarya* isolates to characterize black pod rot resistance. The goals of this study were to: 1) study the genetic variation of the IC3 cacao collection and the *P. megakarya* population, 2) proactively identify potential host resistance sources, and 3) test whether incorporating pathogen genomic variation improves genomic prediction of host resistance.

As one of the two international cacao collections, the IC3 at CATIE in Turrialba, Costa Rica, maintains more than 1,100 cacao clones from Central America, South America, the Caribbean, Asia and Africa, although wild genotypes from the upper and lower Amazon are predominant (Mata-Quirós et al., 2018). The pathogen capture panel was designed to interrogate genes implicated in infection processes, including effector repertoires and enzymatic gene families involved in host tissue colonization. Together, these marker resources enabled genetic characterization of both the host and pathogen populations. Resistance was phenotyped as lesion diameter on unwounded, detached pods four days after zoospore inoculation, using a sparse design comprising 1,238 cacao pods from 350 clones and 58 *P. megakarya* isolates. Detached-pod and leaf-disc tests both correlate with black pod rot field resistance at the level of genotype means (Iwaro et al., 2005; Tahi et al., 2006) and neither consistently outperforms the other.

We used mixed linear models to evaluate how genomic information affected variance component estimates and the prediction of lesion size. The models progressively incorporated genomic relationship matrices for cacao clones and *P. megakarya* isolates, followed by a dual-genome similarity matrix to account for host–pathogen interactions. These analyses were conducted under a sparse screening design, in which the availability of cacao pods largely determined which clones could be evaluated in each bioassay. We further characterized the population structure of both host and pathogen collections, assessed the stability of host performance across pathogen subpopulations, and explored ancestry-based trends in predicted resistance as a hypothesis-generating framework for breeding.

By explicitly testing genomic covariance structures for both host and pathogen, this study evaluates when dual-genome information improves prediction in a sparse perennial-crop disease screen. More broadly, it demonstrates how dual-genome modeling approaches can be implemented in perennial crop pathosystems characterized by complex population structure, sparse testing design, and genetic diversity on both sides of the host-pathogen interaction.

## Methods

### Genetic Material

#### Theobroma cacao

The germplasm collection at the International Cacao Collection (IC3) at CATIE in Turrialba, Costa Rica, was sampled for genotyping and provided pods for disease screening assays. The collection at the “La Montaña” site, located at 602 m above sea level, with a mean annual rainfall of 2,645 mm and a mean temperature of 22.5°C (Mata-Quirós et al., 2017; W. M. Phillips-Mora et al., 2007). The collection was re-established at this site between 2001 and 2005, with three trees per accession, and comprises approximately 1,200 accessions representing diverse cacao genetic backgrounds (Mata-Quirós et al., 2018; W. M. Phillips-Mora et al., 2007). No disease control was carried out other than the removal and disposal of diseased fruits. Naturally pollinated pods approximately 4 months old (unripe) were harvested in the morning, individually wrapped in foam, placed in a shipping box, and sent to the Plant Pathology Department of the University of Florida in Gainesville, Florida, USA, for screening with *Phytophthora* sp. isolates. Pods were imported under APHIS PPQ permit 556-20-330-01055 and 556-23-242-14283.

#### Phytophthora megakarya

The initial study panel comprised 67 *Phytophthora megakarya* isolates originating from Ghana (21), Cameroon (18), and Nigeria (28). Isolates were obtained from collections maintained by the USDA in Beltsville, USA, Gerben Martijn ten Hoopen, and the University of Florida (Supplementary Table 1), and were imported or transferred under APHIS PPQ permits P526P-20-02935 and P526P-21-01779. Isolates originated from black pod-infected cacao or soil samples collected in West Africa. Previously characterized isolates were obtained from collections described by Ali et al. (2016), while additional isolates originated from epidemiological studies of *P. megakarya* in Cameroon (Ndoungué Djeumekop et al., 2021). Nigerian isolates originated from the states of Abia, Cross River, Edo, Ondo, and Osun, whereas Cameroon isolates originated from Bakoa and Kédia (Supplementary Table 1). Species identity was confirmed by PCR using the *P. megakarya*-specific primers and amplification conditions described by Ali et al. (2016).

All isolates were revived by inoculating cacao pod tissue, followed by re-isolation and subsequent storage. Wounded pod-husk inoculation assays were conducted using an adaptation of the methods described by Ali et al. (2017) and Puig et al. (2018). Briefly, cacao pod husks were cut into 6 × 6 cm pieces and placed in sterile plastic containers for incubation. Following lesion development, infected tissue was excised from the advancing lesion margin using a sterile scalpel, and three tissue pieces were transferred to 5% clarified tomato juice (CT) agar (1.5% agar) amended with 5 mg L⁻¹ pimaricin, 250 mg L⁻¹ ampicillin, and 10 mg L⁻¹ rifampicin (PAR), adapted from Ferguson and Jeffers (1999). After hyphal growth was observed, mycelium was transferred to water agar (1.5% agar) for purification, followed by transfer to 10% CT agar (1.5% agar, PAR). For long-term storage, colonized agar plugs (5 × 5 mm) were transferred to autoclaved 8-mL glass vials containing reverse-osmosis water, either with or without sterilized hemp seeds, in duplicate, using a method adapted from Sutton et al. (2015). Alternatively, isolates were maintained on 10% CT agar (1.5% agar, PAR) in the dark at room temperature for short-term storage.

### Genotyping

#### Theobroma cacao

One healthy young leaf was collected from each clone in February 2022 in the IC3 collection in Costa Rica, with the first tree in the three-tree row plot sampled if available. Twelve clones were not sampled because no living trees were available. Leaf samples were collected in paper bags and shipped to Mars Plant Science Center (Davis, CA), with silica gel bags used to capture humidity. BioArk Leaf collection kits (LGC Biosearch Technologies, Teddington, Middlesex, UK) were used to collect eight 5 mm leaf discs from a single leaf for each sample. The leaf disc samples were shipped to LGC for DNA extraction using their sbeadex™ extraction protocol followed by genotyping with SeqSNP^©^ (LGC Biosearch Technologies). SeqSNP^©^ employs an enrichment approach to target specific regions of the genome for sequencing. The resulting sequence data was aligned to the Matina V1 cacao reference (Motamayor et al., 2013) using bwa (Li & Durbin, 2009) and freebayes v1.2.0 (Garrison & Marth, 2012) was used to call variants for all cacao clones. The targeted regions consisted of 10,000 preselected positions, most of which had been previously published (Allegre et al., 2012; Jarquín et al., 2014; Livingstone et al., 2015, 2017; Romero Navarro et al., 2017). The panel contained three sets of markers: 1) 8,635 markers that were evenly distributed across the genome, 2) 864 markers within nucleotide-binding and leucine-rich repeat genes that were identified in the ‘Criollo’ genome v2 (Argout et al., 2017) using the NLRannotator pipeline v.0.7 (Steuernagel et al., 2020), and 3) 501 markers associated with genes or quantitative trait loci identified in various other studies. The resulting VCF-file was processed using a customized script in R 4.3.2 (R Core Team, 2023) and RStudio v.2025.5.0.496 (Posit team, 2025). The clone names were standardized and harmonized, and biallelic SNPs were retained and genotypes were encoded as additive dosages of 0, 1, or 2 copies of the alternate allele using the package SNPassoc 2.1-0 (Moreno et al., 2022). If multiple samples of the same clone were present, similarity was verified using the correlation of marker calls. For samples showing a correlation below 95% with the same clone and greater than 98% with one or more other clones, the location in the orchard was added to the clone’s name.

An assignment test was applied to infer population membership and admixed ancestry (hybrids or ancestral forms) of the clones in the 11 reference groups, using a model-based clustering method implemented in the software program STRUCTURE (Pritchard et al., 2000). The analysis used selected SNPs and reference accessions representing ‘Amelonado’, ‘Bolivian’, ‘Contamana’, ‘Criollo’, ‘Curaray’, ‘Guiana’, ‘Iquitos’, ‘Maranon’, ‘Nacional’, ‘Nanay’, and ‘Purus’. STRUCTURE was run under the admixture model with correlated allele frequencies, using a burn-in of 100,000 iterations followed by 200,000 MCMC iterations. Ten replicate runs were performed with the number of clusters (K-value) set to 11, assuming that each of 11 reference groups may have contributed to the clones. For each clone, the STRUCTURE membership coefficients (Q) were interpreted as ancestry proportions. Clones with ≥75% membership in their predominant reference group were classified as members of that group; clones below this threshold were classified as admixed and named according to the ancestry component with the highest Q value.

#### Phytophthora megakarya

*P. megakarya* isolates were grown on 10% CT plates at room temperature for 7 days in the dark. Three agar plugs were transferred to plates, covered with 20 mL of 10% liquid CT, and grown at 25°C in the dark for 7 days. Mycelia were washed, harvested, and concentrated using a vacuum pump prior to storing at -80°C and freeze-drying for 24 hours using a FreeZone 2.5 Liter -50C Benchtop Freeze Dryer (Labconco Corporation, Kansas City, MO, USA). LGC Biosearch Technologies Inc. (Hoddesdon, UK) performed DNA extractions and library preparations using their high-throughput workflow with proprietary chemistry. DNA was sheared to a mean fragment length of 500 bp, fragments were end-repaired and A-tailed, followed by incorporation of unique dual-indexed Illumina adaptors and PCR enrichment, equimolar pooling, and library preparation. Similar to cacao, enrichment was employed prior to sequencing to focus on virulence-related genes selected from the *P. megakarya* and *P. palmivora* genomes, with annotations published by Morales-Cruz et al. (2020). The target panel comprised gene families critical to the infection process, e.g., proteases, CAZymes, detoxification genes, and effectors (Supplementary Table 2). LGC Biosearch Technologies Inc. (Hoddesdon, UK) designed and synthesized the biotinylated probes used to enrich the targeted regions, providing a 1x probe tiling across the targets. Probes were evaluated for GC content, homopolymers, and uniqueness within the genome and filtered as needed. The final panel consisted of 54,046 probes, each 120 bp in length. These probes were designed to target 3,505 *P. megakarya* genes, 2,774 SNP locations previously identified using genotype-by-sequencing (unpublished), and 2,381 *P. palmivora* genes (Supplementary Table 2). The DNA libraries were hybridized to the capture-Seq probes and sequenced on a NovaSeq S4 flow cell (2 x 150 bp). The samples were multiplexed, and demultiplexing was done by LGC Biosearch Technologies Inc. (Hoddesdon, UK).

The sequencing data were checked for quality using FastQC v0.12.1 (Andrews, 2010), and MultiQC 1.24.1 (Ewels et al., 2016) was used to compile a summary report. Trimming was done using fastp 0.22.0 (Chen et al., 2018) with default settings. Filtered high-quality reads were mapped to the *P. megakarya* GH34 genome (Morales-Cruz et al., 2020) using BWA-MEM 0.7.17 (Li, 2013) with the “-M” flag to mark shorter split hits as secondary. The aligned SAM files were processed with SAMtools 1.19 (Danecek et al., 2021) to convert them to BAM files, sort, fix mate pair information, remove PCR duplicates, filter for quality (MAPQ ≥ 20), and obtain overall mapping statistics. SNP calling was performed per sample and merged using GATK v.4.4 (Van der Auwera & O’Connor, 2020) *HaplotypeCaller* with ploidy set to 2 and *CombineGVCFs*. SNPs were extracted and filtered using the recommended filtering criteria by GATK for read and mapping quality, as determined by thresholds for quality by depth (QD <= 2.0), strand bias (FisherStrand (FS) > 60.0), and root mean square mapping quality over all the reads (MQ < 40.0), and filtered for the target regions. Summary statistics were generated using rtg-tools 3.7.1 (Cleary et al., 2015) and bcftools 1.22 (Danecek et al., 2021). The resulting VCF file was analyzed in R 4.3.2 (R Core Team, 2023) using the vcfR package v1.15.0 (Knaus & Grünwald, 2017) for extraction, filtering, and conversion, and the ggplot2 package 3.5.0 (Wickham, 2016) for visualization. Because sequencing depth varied among isolates, genotype calls in the lowest and highest 10% of the depth distribution for each sample were excluded before downstream analysis; the corresponding depth thresholds ranged from 27–46× for the lower cutoff and 544–1,577× for the upper cutoff. Biallelic SNPs were retained and encoded as additive genotype dosages (0, 1, or 2), leaving 131,162 SNPs across 67 isolates.

### Population structure and genomic relationship matrices

The final marker matrices from genotyping were processed using ASRgenomics 1.1.4 (Gezan et al., 2022) for quality control (QC) and the construction of relationship matrices. Relationship matrices were calculated using Van Raden’s method (VanRaden, 2008) to convert the n × m marker matrices into n × n genomic relationship matrices (GRM) (Amadeu et al., 2023), where n is the number of individuals and m the number of markers. Of the 1,208 cacao samples submitted for genotyping, 28 samples were duplicates (r_i,j_=1). The samples with fewer missing values across the marker set were kept for analysis. Marker filtering was performed using thresholds for missing values within individuals (≥0.5), or within markers (≥0.5), and rare markers with minor allele frequency (≤0.05) were removed. This filtering process removed four individuals and 251 markers from the cacao population, leaving 1,176 individuals x 8,593 markers. The GRM was computed, with missing data imputed using marker-specific average frequencies. Kinship diagnostics used to detect inbreeding and anomalies in relatedness estimates (F_i_ ≥ 1.9 and ≤ 0.5) identified 159 individuals that were removed prior to a second round of kinship filtering, that resulted in the removal of an additional 19 individuals. A final kinship filtering step was applied to remove duplicates (r_ij_ > 0.98), resulting in a cacao GRM (GRM_c_) based on 947 individuals and 8,572 markers. The names of clones identified as duplicates were updated to match the remaining clone in the analysis.

Relationship matrices for the 67 *P. megakarya* isolates were created following marker filtering using thresholds for missing values within individuals (≥0.5), or within markers (≥0.25), and the minimum minor allele frequency (≤0.05). These filtering steps removed 67,227 markers. An additional 104 homozygous SNPs across the population were also removed (MAF = 1), resulting in a matrix of 67 individuals x 63,831 markers. Base R and the software packages ‘factoextra’ (Kassambara & Mundt, 2016) and ‘FactoMineR’ (Husson et al., 2006) were used to investigate and identify the population structure in both *Theobroma cacao* and *P. megakarya*. The extreme population structure of the *P. megakarya* isolates was evident when the first two components of a principal component analysis (PCA) using filtered markers were examined (Supplementary Figure 1). The number of subpopulations for *P. megakarya* was defined using PCA and hierarchical clustering. For each subpopulation, the marker matrix was extracted, marker filtering was applied, and a GRM was computed.

VanRaden genomic relationships are derived from allele-frequency-centered marker dosages, and their scale depends on the marker-frequency distribution used in the population. These assumptions are not met in a predominantly clonal organism like P. megakarya. The mean diagonals of the unscaled subpopulation matrices were accordingly 0.19, 0.41 and 0.39 for subpopulations A, B and C, compared to 1.15 for the cacao matrix. Because a variance component estimated against a relationship matrix K is expressed on the scale of K, each matrix was divided by D = mean(diag(K)) - mean(K) (Legarra, 2016) so that its variance component equals the variance of the genetic values it describes. Each *P. megakarya* subpopulation matrix was scaled by its own D before each subpopulation-specific GRM was combined into a block-diagonal multi-population isolate similarity matrix, so that a single isolate variance component applies on a common scale across the three subpopulations. All variance ratios reported below are therefore comparable among models and among pathogen subpopulations. The final GRM for isolates was defined as the block-diagonal matrix GRM_I_:

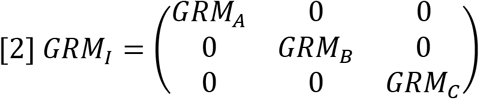

The final pathogen GRM had a block-diagonal structure to model genomic covariance among isolates within each subpopulation, whereas covariance between subpopulations was set to zero. Broad differences among subpopulations were therefore modeled separately as fixed effects.

The following packages were used for visualizations in R: ggplot2 3.5.0 (Wickham, 2016), ggrepel 0.9.5 (Slowikowski, 2016), ggsci 3.0.0 (Xiao, 2016), ggpubr 0.6.0 (Kassambara, 2016), rcartocolor 2.1.1 (Nowosad, 2017), scales 1.3.0 (Wickham et al., 2011), ggvenn 0.1.19 (Yan, 2021), and superheat 0.1.0 (Barter & Yu, 2018).

### Disease screening

Variation in the timing of pod availability across CATIE’s IC3 germplasm collection prevented the use of balanced experimental designs. The germplasm collection was sampled as unripe 4-month-old pods were available. Fruit ripening stages were assessed visually based on genotype-specific changes in pod color associated with maturation (W. Phillips-Mora et al., 2013). A total of 31 shipments from the CATIE germplasm collection to the University of Florida defined different bioassays, with each bioassay containing up to 100 pods. As many as five isolates were used to inoculate the clones in a single bioassay. A total of 1,971 pods, representing 563 clones, were inoculated with one or several of the 67 *P. megakarya* isolates between May 2021 and May 2023. After the first year of screening, a selection of clones was made from the clones phenotyped to date to represent the collection’s genetic diversity and facilitate replication of clone-by-isolate combinations. POUND-7 served as control clone and was tested in 22 out of 31 bioassays, because pods were readily available from a planting adjacent to the IC3 collection and because it has demonstrated resistance to black pod rot caused by *P. palmivora* (Brown et al., 2007; W. Phillips-Mora & Galindo, 1989). *P. megakarya* GH34 served as a control isolate because its published genome annotation was used as the reference for probe design (Morales-Cruz et al., 2020), and its aggressiveness on cacao pods had previously been characterized (Ali et al., 2016; Puig et al., 2018). The isolate was grown and prepared for every bioassay and used if one of the selected isolates did not produce enough zoospores for inoculation (11 of 31 bioassays).

Initial experimentation led to the development of a bioassay protocol, which has been deposited in the protocols.io repository (Gitto et al., 2026). This artificial inoculation technique is based on the paper disc method developed by Phillips-Mora and Galindo (1989). The bioassay was designed to yield six observations per pod. Pods were prepared for inoculation by cleaning the surface with distilled water. While pods from different clones were harvested at the same time, differences in shipping times cause the time between harvest and inoculation to vary from 4 to 13 days (average 8 days). Damaged pods or pods with obvious disease symptoms that developed during shipping were discarded. The plates used to create the zoospore solutions were prepared 1 week in advance of inoculation. Colonized agar mycelium chunks from up to five isolates were placed on two 10% CT plates per isolate and prepared as described in the protocol. For inoculations, zoospore solutions were prepared at a concentration of 50,000 spores per ml, and 1 cm² pieces of tissue paper were placed on the pod prior to pipetting 30 μl of the solution into six evenly spaced positions on the pod surface. Twelve inoculated pods were placed in each incubation box, and up to six boxes were placed in two inoculation chambers (Percival Scientific, model I36NL) to maintain a temperature of 25°C at a humidity of approximately 100%. Four days after inoculation, vertical and horizontal measurements were taken for each of the six inoculation sites per pod using calipers. Lesions were measured on both the vertical and horizontal axes, and the average of both lesion measurements was calculated per inoculation site. All analyses of lesion size were then performed using the average of the six mean lesion measurements per pod. Lesions less than 15 mm were within the initial area of the tissue paper-based inoculation. A response only in this area was considered resistant, because the lesion did not spread beyond the inoculation area. Lesion measurements below 30 mm were considered moderately resistant, whereas measurements above 30 mm indicated susceptibility.

### Statistical models

Data were analyzed using a set of linear mixed models to produce variance component estimates and provide lesion-size predictions for all clones, isolates, and clone-by-isolate interactions observed in the bioassays, as well as tests of significance for model effects. Covariance structures associated with the random effects in the mixed models were used to compare models that used traditional design matrices to account for bioassay effects, as well as marker-derived relationship matrices for cacao clones (GRM_C_), *P. megakarya* isolates (GRM_I_), and clone-by-isolate interactions. Variance ratios for host, pathogen, and host-by-pathogen effects and type-B correlations for host and pathogen were derived from the fitted variance components from the general linear mixed model, presented in matrix notation following Mrode and Thompson (2005) as:

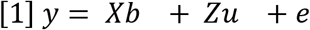

where: **y** refers to phenotypic observations (mean lesion diameter per pod in mm); **b** is the vector of fixed *P. megakarya* subpopulation effects; **u** is the vector of random effects associated with the bioassay, cacao clone, *P. megakarya* isolate, and the cacao clone by *P. megakarya* isolate interaction; **X** is the design matrix relating each observation to the **b** fixed effects; **Z** is the design matrix relating the **y** observations to the **u** random effects; and **e** is the vector of residuals or random errors associated with each observation. Random effects for clone, isolate, clone-by-isolate interaction, and bioassay were assumed to be normally distributed, with σ^2^_Clone_, σ^2^_Isolate_, σ^2^_Clone:Isolate,_ and σ^2^_Bioassay_ defining the variance components associated with each random effect, respectively. Separate error variances were estimated for each *P. megakarya* subpopulation to allow for heteroskedasticity of the error variances, and the average of these error variances (σ^2^_E_) was used to estimate genetic parameter estimates.

Six mixed models differing in their covariance structures for host, pathogen, and host-by-pathogen effects were fitted to the same phenotype dataset (Table 1). Model fit was compared using the REML log-likelihood (logL) and Akaike’s information criterion, AIC = −2logL + 2k, where k is the number of estimated variance parameters. All models were fitted to the same records with the same fixed effect (subpopulation), so their REML log-likelihoods are comparable. Several pairs of models are not nested, so models were ranked by AIC; a lower AIC indicates a better fit. Nested comparisons of single variance components used likelihood ratio tests, with the null distribution taken as a 50:50 mixture of χ²₀ and χ²₁ because the variance is tested at its boundary (Self & Liang, 1987).

**Table 1:** Comparisons of genetic parameter estimates of all models tested in this study. The estimations of the percentage of phenotypic variance (ratios on VP = clone + isolate + interaction + residual, bioassay is fitted but excluded from V_P) with standard errors explained by genetic effects for cacao clones (*Ĥ*^2^), *P. megakarya* isolates, their interaction, and the residual error term (*e*^2^), along with estimates of genetic correlations with standard errors for clones *r*_gC_ and isolates *r*_gI_ are given. Those were obtained from linear models incorporating the factors clone and isolate as either unrelated and independent design effects or as correlated effects through marker-derived relationship matrices (GRM) for cacao clones and *P. megakarya* isolates. A dual-genome similarity matrix (GGSM) was used to model interactions. Model 6 fits subpopulation-specific genomic variance components via a heterogeneous correlation structure for clones and subpopulation-specific isolate effects. The model also provides a genomic correlation between subpopulation (cor.). The subpopulations are A, B, and C. The cacao ratio is a genomic broad-sense heritability; the isolate and interaction ratios are proportions of phenotypic variance. Phenotypic variance is defined excluding the bioassay variance. Each relationship matrix was scaled by *D* = *mean*[*diag*(*K*)] − *mean*(*K*), which was 1.136 for cacao and 0.192, 0.413, and 0.391 for pathogen subpopulations A, B, and C. logL, REML log-likelihood; AIC, Akaike information criterion (−2logL + 2k, where k is the number of estimated variance parameters: 7 for Models 1–5 and 11 for Model 6). Lower AIC indicates better fit.

| # | Model structure | $\widehat{H}_C^2$ | isolate | clone: isolate | e² | $r_{gC}$ | $r_{gI}$ | logL | AIC |
|---|---|---|---|---|---|---|---|---|---|
| 1 | C, I, C:I | 0.199 ± 0.028 | 0.362 ± 0.055 | 0.016 ± 0.026 | 0.423 ± 0.046 | 0.925 ± 0.111 | 0.957 ± 0.065 | −3768.083 | 7550.17 |
| 2 | GRM<sub>C</sub>, I, GRM<sub>C</sub>:I | 0.267 ± 0.039 | 0.344 ± 0.053 | 0.017 ± 0.014 | 0.373 ± 0.039 | 0.941 ± 0.047 | 0.953 ± 0.038 | −3744.866 | **7503.73** |
| 3 | C, GRM<sub>I</sub>, C:GRM<sub>I</sub> | 0.195 ± 0.028 | 0.378 ± 0.057 | 0.012 ± 0.025 | 0.416 ± 0.047 | 0.944 ± 0.113 | 0.970 ± 0.061 | −3769.399 | 7552.80 |
| 4 | GRM<sub>C</sub>, GRM<sub>I</sub>, C:I | 0.258 ± 0.039 | 0.361 ± 0.056 | bound | 0.381 ± 0.039 | 1 (bound) | 1 (bound) | −3747.313 | 7508.63 |
| 5 | GRM<sub>C</sub>, GRM<sub>I</sub>, GGSM<sub>CI</sub> | 0.258 ± 0.039 | 0.360 ± 0.056 | 0.008 ± 0.014 | 0.374 ± 0.040 | 0.971 ± 0.053 | 0.979 ± 0.038 | −3747.170 | 7508.34 |
| 6 | *corh*(subpop: GRMc), *at*(subpop):GRM<sub>I</sub> | A 0.298 ± 0.059/<br>B 0.191 ± 0.071<br>C 0.328 ± 0.061 | A 0.396 ± 0.092<br>B 0.489 ± 0.124<br>C 0.262 ± 0.079 | n/a | A 0.307 ± 0.052<br>B 0.320 ± 0.086<br>C 0.410 ± 0.058 | n/a | n/a | −3734.655 | **7491.31** |

Models 2–5 progressively replaced identity covariance structures with marker-derived relationship matrices for cacao clones, *P. megakarya* isolates, or clone-by-isolate combinations. Model 1 used traditional design matrices for clone, isolate, and interaction effects, assuming no covariance or relatedness among cacao varieties or *P. megakarya* isolates. In Model 2, GRM_C_ and a GRM_C_-by-isolate design matrix accounting for clone-by-isolate interaction effects replaced the traditional design matrices that were used for Model 1. In Model 3, GRM_I_ and GRM_I_-by-clone design matrices replaced the Model 1 design matrices. In Model 4, GRM_C_ and GRM_I_ replaced their respective Model 1 design matrices. For the replacement of the clone-by-isolate interaction design matrix, the cacao GRM_C_ and the *P. megakarya* GRM_I_ were combined to form a covariance matrix, which was incorporated into a reaction norm model (Jarquín et al., 2014), to generate a dual-genome similarity matrix (GGSM_CI_) (Hudson et al., 2024). Model 5 incorporated the GGSM_CI_ rather than the identity matrix to account for similarities among clone-by-isolate interactions. The significance of clone, isolate, and interaction effects were evaluated with likelihood ratio test that utilized Model 2 as the complete model for comparisons with reduced models that excluded the random effects under test. A final Model 6 used GRM_C_ with a heterogeneous correlation structure (*corh()*) for isolate subpopulations that allowed for separate genomic variances for each isolate subpopulation. Additionally, subpopulation-specific isolate effects were included using the *at()* directive of ASReml. A full unstructured covariance model (*corgh()*) for clone genomic effects was initially evaluated but converged with the correlation between subpopulations A and B at its boundary (1.00), so the heterogeneous model with a single correlation (corh()) is reported. Predictions for cacao clones, *P. megakarya* isolates, and clone-by-isolate effects are derived from Model 2, with subpopulation-specific clone predictions from Model 6 to provide comparisons among cacao clones and *P. megakarya* isolates and subpopulations, respectively (Supplementary Tables 5 and 6). Model 6 had the lowest AIC, but its additional parameters described subpopulation-specific variance heterogeneity rather than a different overall ranking of host clones as the estimated genomic correlation among subpopulations was high (0.86). We therefore used Model 2, which provides a single genome-wide host effect, for cross-validation and overall clone prediction.

### Genetic parameter estimates

Genetic parameter estimates are provided for each model. For each model, the proportion of phenotypic variance attributable to the cacao clone component (*Ĥ*^2^) was calculated as the ratio of genetic variance associated with variation among clones to the total phenotypic variance:

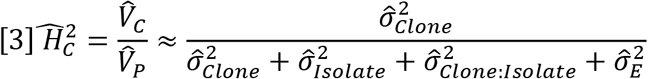

with variance components defined above, *V̂*_P_ as defined is the sum of the clone, isolate, interaction and residual variances, not the total observed phenotypic variance, since the bioassay effect is still fitted but not counted. The bioassay variance was 49–57 mm² across models, about 16% of the total variance (Supplementary Table 7). *V̂*_C_ is the genotypic variance of cacao clones. Analogous variance ratios were calculated for isolate (*V̂*_I_) and clone-by-isolate effects (*V̂*_CI_) to permit direct comparison of fitted variance components on the same phenotypic scale:

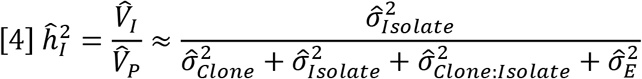

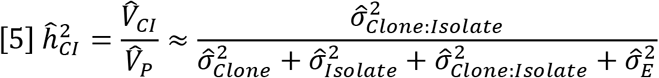

Type-B correlations were calculated as variance-ratio summaries of the relative magnitude of clone-by-isolate interaction (Burdon, 1977) following equations 6 and 7.

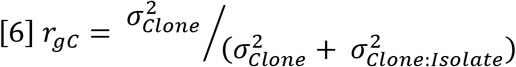

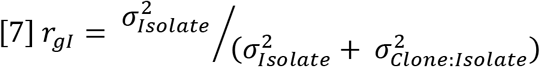

Both quantities are bounded functions of the interaction variance and approach unity as the interaction variance approaches zero, so they are reported as descriptions of the fitted variance partition typically used to indicate the stability of lesion size predictions across isolates (*r_gC_*) or clones (*r_gI_*).

### Cross-validation

The ability of Model 2 to predict phenotypically unobserved clones within the IC3 population was evaluated using clone-wise 10-fold cross-validation. The 350 phenotyped clones were assigned at random to ten folds. For each validation fold, all lesion observations from the corresponding clones were masked and Model 2 was refitted to the remaining records, so validation clones were predicted solely through genomic relationships with phenotyped training clones. Predictive ability was the Pearson correlation between the predicted genomic values of the masked clones and their observed clone means. Before averaging, each pod was adjusted for the subpopulation, isolate, and bioassay effects estimated from Model 2. To obtain prediction accuracy, we divided predictive ability by the square root of the mean heritability of clone means:

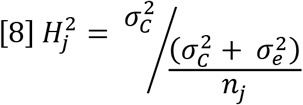

where *n*_j_ is the number of pods of clone j. We repeated the procedure with five random partitions. To test whether the pathogen GRM improves prediction, we evaluated Model 4 on the same partitions and against the same adjusted clone means and compared the two models within each partition.

### Genetic-group ancestry and predicted resistance

To relate predicted resistance to the genetic groups in the collection, we regressed the Model 2 GEBVs of the 350 phenotyped clones on their STRUCTURE ancestry proportions (K = 11) in a single multiple regression. Ancestry proportions sum to one, and ‘Amelonado’, the most common group, served as the reference. With ‘Amelonado’ omitted as the reference component, each coefficient represents the expected difference in GEBV associated with a one-unit increase in the indicated ancestry proportion relative to ‘Amelonado’. Collinearity was checked using variance inflation factors. Unphenotyped clones were excluded, because their GEBVs were derived entirely from phenotyped relatives. Because GEBVs are shrunken and mutually correlated through a GRM derived from the same marker panel that informs ancestry, inferential statistics from this regression are not independent of the genomic model. The analysis was therefore treated as exploratory and used to identify ancestry-associated trends rather than estimate causal genetic-group effects.

## Results

### One-third of the IC3 cacao collection shows high ancestry membership to seven genetic groups of cacao

All living clones of the CATIE IC3 collection were genotyped with a 10K SNP marker panel. After calling SNPs, cleaning for SNP quality, inbreeding, and duplications, 947 clones and 8,572 markers remained. The clones were assigned to eleven genetic reference groups described in cacao (Motamayor et al., 2008; Zhang et al., 2012) (Figure 1, Supplementary Table 3). 300 clones were assigned as members of one of seven genetic groups (≥75% contribution), while 647 clones were considered admixed (<75%). The genetic groups ‘Contamana’, ‘Criollo’, and ‘Curaray’ were represented only by admixed clones, while the ‘Bolivian’ genetic group was present, but did not strongly contribute to the examined clones. The genetic group ‘Amelonado’ contributed the most to the IC3 collection, both in terms of members (13.8%) and hybrids (34.4%), while ‘Curaray’ showed the lowest contribution as a genetic group (0.7%) (Figure 2C). The average proportion of ‘Amelonado’ across clones in the whole population was 39.6%, followed by ‘Criollo’ and ‘Nacional’ with an average proportion of 10% each. There was an overlap of 131 clones with the population described in Motamayor et al. (2008), with 81% of these clones assigned to the same genetic group (Supplementary Table 3). The relationships among all clones were visualized using PCA, with the first two components accounting for 25.2% of the total genetic variation (Figure 2A and 2B). Ancestry-related genetic structure was evident in the genome-wide SNP-based PCA among clones assigned as genetic group members (Figure 2A). Clones with ≥75% ‘Amelonado’ ancestry occupied a broad and partially structured region of PCA space across PC1–PC3, indicating substantial within-group diversity. ‘Guiana’ members formed a compact region at high PC2 scores, while ‘Maranon’ and ‘Nanay’ also occupied comparatively high PC2 positions but were distinguished with additional components (Supplementary Figure 2A and 2B). ‘Iquitos’ and ‘Purus’ occurred near one another along PC1 but separated more clearly on PC3 and PC4. ‘Guiana’ remained distinct across the first four principal components, while the numerous admixed clones occupied intermediate regions among the high-membership groups.

**Figure 1:**
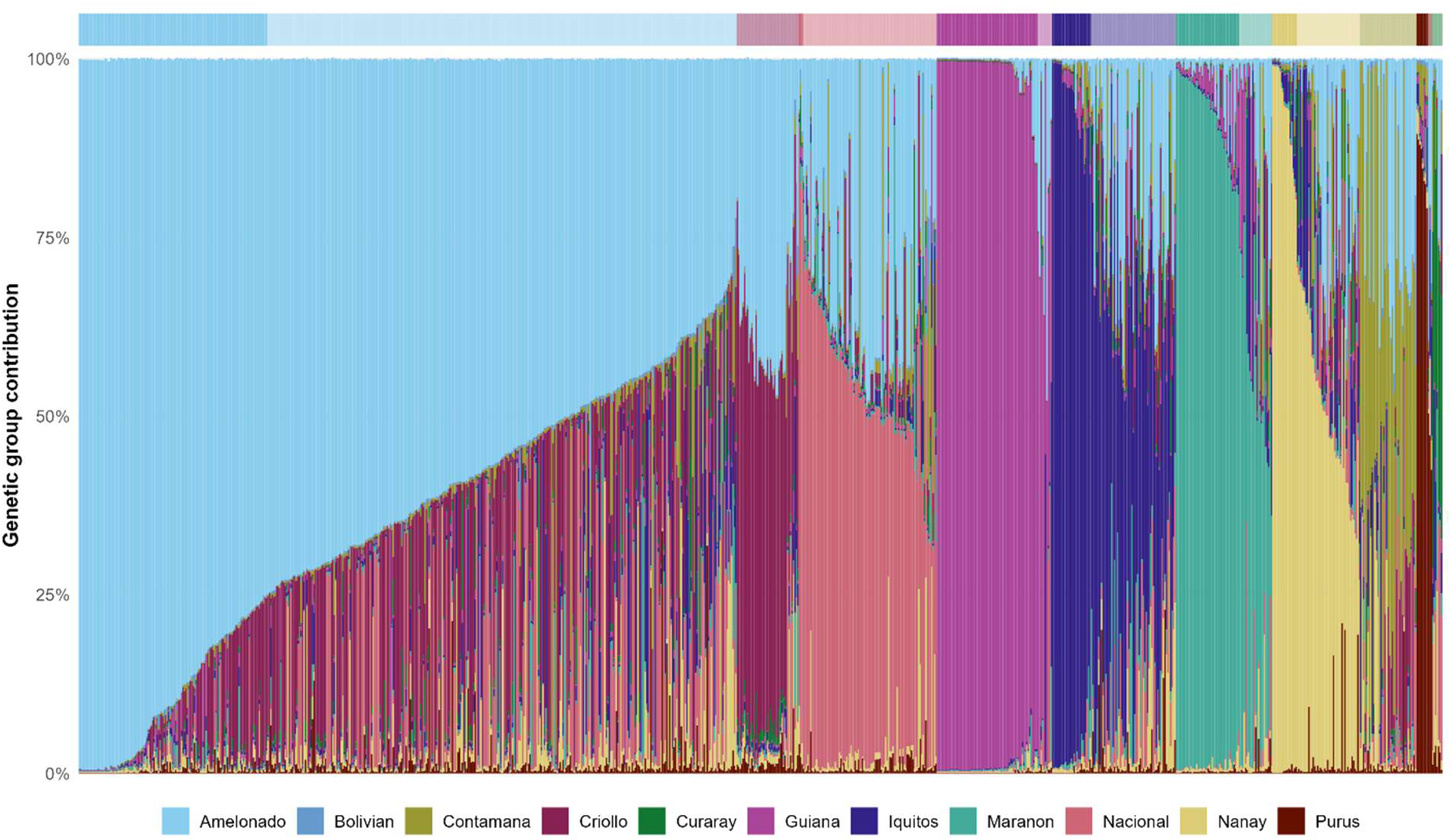
STRUCTURE analysis showing the genetic group proportion relative to the eleven genetic groups of *Theobroma cacao* for all clones from the IC3 cacao collection included in the trial. Ten genetic groups were described by Motamayor et al. (2008), and the genetic group ‘Bolivian’ was described by Zhang et al. (2012). The color bar above the plot indicates whether the clone is considered a full member of a genetic group (≥75% membership; colored according to the legend) or an admixed clone (25-75%, lighter color corresponding to the genetic group with the highest contribution).

**Figure 2:**
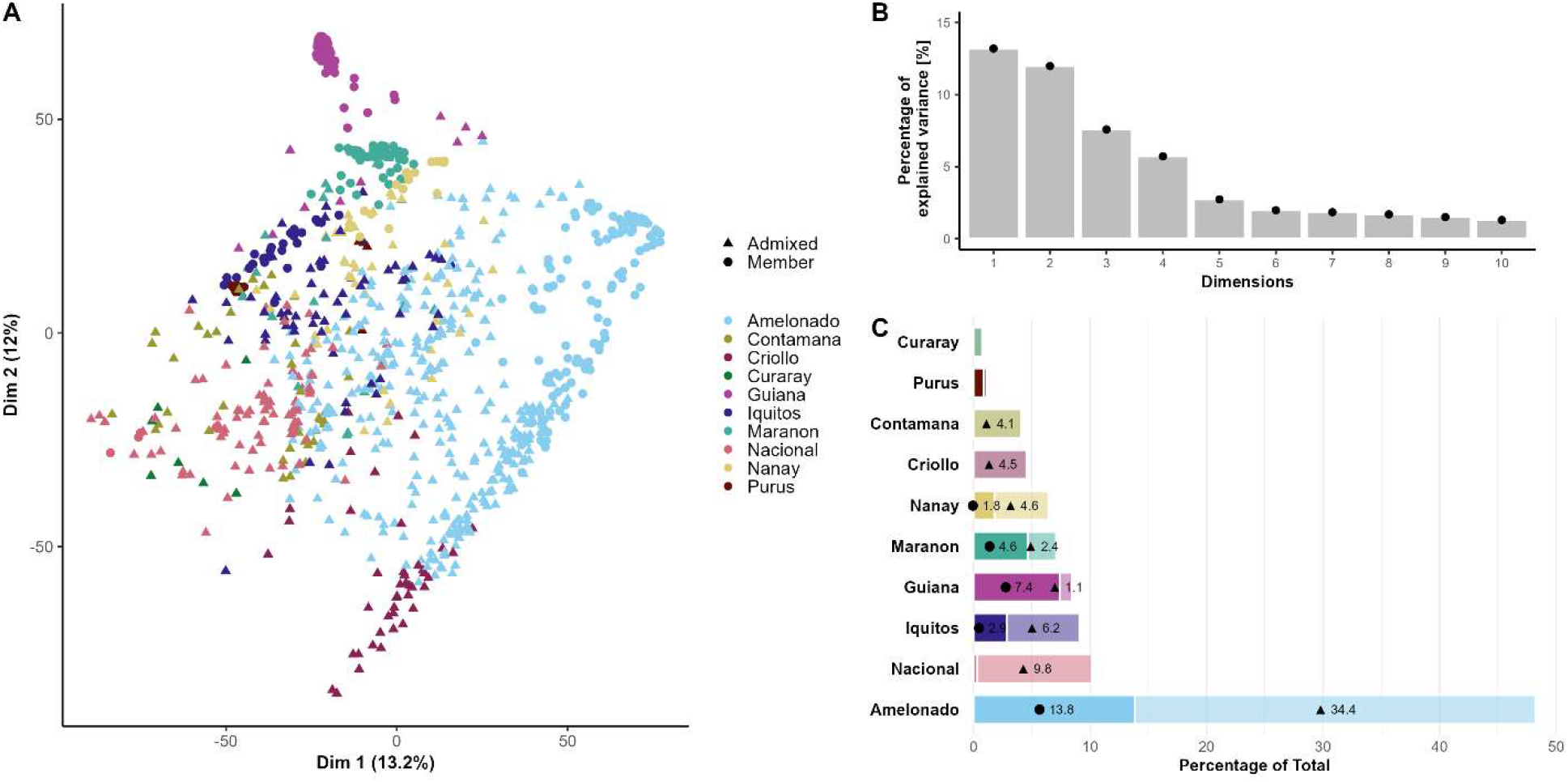
Population structure and composition of 947 clones of the IC3 cacao collection. **(A)** Principal component analysis (PCA) based on 8,572 SNP markers. Each point represents an individual clone and is colored according to its predominant genetic-group ancestry based on STRUCTURE analysis. Clones with ≥75% membership in their predominant genetic group were classified as members and are shown as circles, whereas clones with <75% membership were classified as admixed and are shown as triangles. **(B)** Percentage of genetic variation explained by the first ten principal components of the SNP-based PCA, **(C)** Representation of the major cacao genetic groups within the IC3 collection, based on STRUCTURE ancestry assignments. Contributions from clones with >75% membership and admixed clones are indicated by circles and triangles, respectively. Genetic-group nomenclature follows Motamayor et al. (2008).

### Three subpopulations were identified in the *P. megakarya* isolates causing black pod

The 67 *P. megakarya* isolates were genotyped using a customized capture-sequencing approach to examine population structure. The main gene families targeted by this approach were effectors (43%), especially secreted RxLRs (36%), proteases (18%), detoxification genes (15%), and CAZymes (13%) (Supplementary Table 2). The rate of duplicated reads ranged from 37.0% to 62.5% with a median of 48.1%. Over 99% of the reads mapped to the *P. megakarya* GH34 genome (Morales-Cruz et al., 2020). PCA of 62,203 filtered SNPs revealed a pronounced separation of isolates into three tight clusters, with PC1 explaining 78.9% of the variance (Supplementary Figure 1). This primary division was used as the first step in defining broad genomic groups, which were then examined separately using PCA and relationship matrices (Supplementary Figures 3 and 4). Within the western group (including isolates from Ghana and Nigeria), additional structure remained evident (PC1 = 48.6%), motivating a further subdivision before construction of within- group GRMs. Additionally, two isolates, “PH009” and “PH021” from Nigeria, were genetically distinct from both subpopulations. The second subpopulation, comprising all isolates from Cameroon and some from Nigeria (Supplementary Figure 4), also separated into two clusters, with one cluster comprising three isolates from Nigeria and one isolate from Cameroon (PC1 = 68.5%). Another Cameroon isolate, “NRRL_64259,” was genetically distinct from both clusters. Seven isolates fell outside the three major genomic groups and were excluded from analyses requiring the block-diagonal pathogen GRM because the small number of outlying isolates did not support estimation of additional group-specific covariance structures. Their exclusion reflects a modeling constraint rather than evidence that these isolates are biologically unrepresentative. The subpopulations were named from West to East. The first subpopulation comprises all isolates from Ghana and two from Nigeria (“A”); the second comprises only isolates from Nigeria (“B”); and the third comprises isolates from Cameroon and five isolates from Nigeria (“C”) (Supplementary Figure 5A-C). After splitting the population into three subpopulations, the marker matrix was extracted per subpopulation. Marker matrix A contained 23 isolates and 11,753 SNPs, out of which 10,871 SNPs (91.6%) were fixed heterozygous. Similarly, marker matrix B contained 14 isolates and 11,614 SNPs, with 10,659 SNPs (91.8%) fixed heterozygous markers. Lastly, marker matrix C contained 21 isolates and 34,519 SNPs, of which 84.6% (29,210 SNPs) were fixed heterozygous. Sites fixed as heterozygous within a subpopulation contain no among-isolate variation within that group and were therefore excluded before construction of the within-subpopulation GRM. The removal of those SNPs in one subpopulation did not preclude finding SNPs in the other two subpopulations. The final marker matrices contained 882, 995, and 5,309 SNPs for the subpopulations A, B, and C, respectively.

Within subpopulations, SNPs covered 4% (A), 6% (B) and 23% (C) of the targeted gene space (Supplementary Table 4). Effectors made up 52.8% of the SNP-containing genes in subpopulation C, against 36.5% in A and 29.3% in B, whereas proteases were relatively more frequent in A and B (28.5% and 30.3%) than in C (17.8%). Most SNPs were specific to one subpopulation (5,258 in C, 765 in B and 723 in A); only 6 SNPs and 27 genes, five of them secreted RxLR effectors, carried variation in all three (Figure 3).

**Figure 3:**
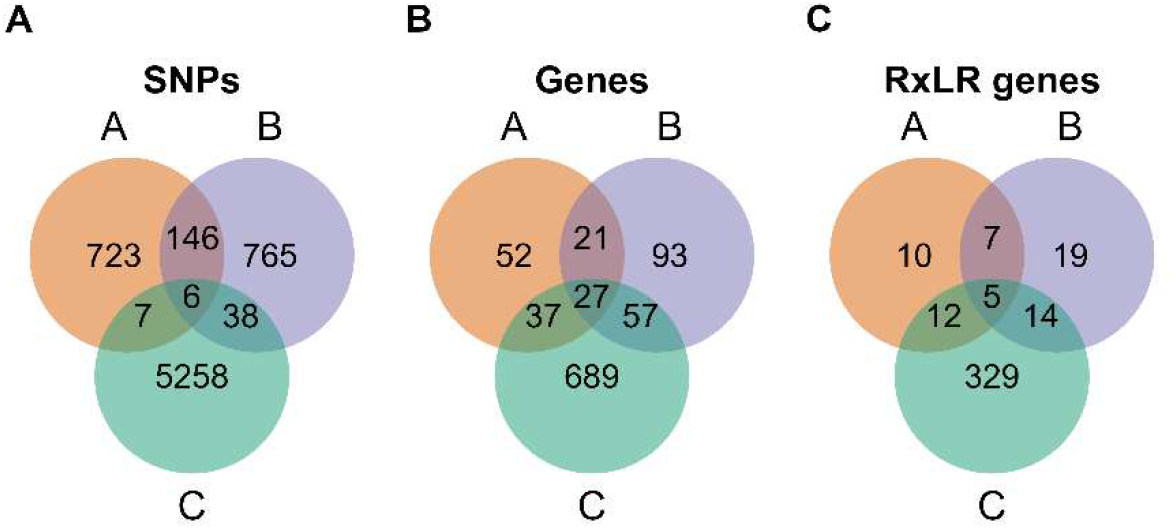
Venn diagram of the total number of SNPs (A), genes with SNPs (B), and RxLR genes with SNPs (C) unique and shared among *P. megakarya* isolates from the subpopulations A, B, and C.

### Variation in lesion size reflects diverse cacao and pathogen responses

Pods that developed disease symptoms unrelated to the controlled inoculations or where physical damage became evident between inoculation and assessment were excluded (19%). Two isolates (N174 and N339) produced no visible infection on >80% of the clones they inoculated and were excluded from the primary quantitative lesion-size analysis (Supplementary Table S1), resulting in a dataset of 1,508 pods from 435 cacao clones inoculated with one or more of the 64 *P. megakarya* isolates. Exclusion of the seven isolates not represented in the three modeled genomic subpopulations resulted in the final analysis dataset of 1,397 pods from 404 clones inoculated with 58 isolates and assessed for lesion size in 30 independent bioassays between 2021 and 2023. Due to the availability of pods across bioassays and by design, 5.1% (1,184) of the 23,432 potential clone-by-isolate combinations were tested. *P. megakarya* isolates were used to inoculate 4 to 78 different clones (median = 18.5, Figure 4A) apart from the reference isolate GH34, which was used to inoculate 227 pods (16%) from 88 different clones (22%). Across the 1,653 isolate pairs, the median number of clones tested with both members of the pair was zero: 51% of pairs shared no clone and 87% shared two or fewer. Each clone was inoculated with 1 to 17 isolates (median = 3), except for the control clone POUND-7, which was inoculated with 18 different isolates (31%) on 26 independent pods (Figure 4B). The lesion diameter per pod (4 DAI) ranged from 0 mm to 70.8 mm, with an overall average of 36.7 mm (Figure 4C). The histograms by subpopulation showed that we collected more data for subpopulation A because it contains the highest number of isolates, and we used isolate GH34 across bioassays. Subpopulation B was the smallest (14 isolates) and had the fewest total data points. Lesion-size distributions were mildly shifted toward larger lesions, particularly for isolates from subpopulation A. The data collected on isolate GH34 showed a diverse response among the 88 inoculated clones: although 83% were susceptible (>30 mm), only 17% (15) were moderately resistant, and no clones were resistant. The isolates tested on the reference clone POUND-7 showed a range of virulence levels. POUND-7 was susceptible to seven isolates (39%), six of them from subpopulation A, moderately resistant to six isolates, and resistant to five isolates; eight of the latter eleven isolates are from subpopulation C. In lieu of analyzing trends from observations on the whole dataset (Supplementary Figure 6, Supplementary Figure 7A-D), mixed models were used to draw conclusions across our sparse experimental design.

**Figure 4:**
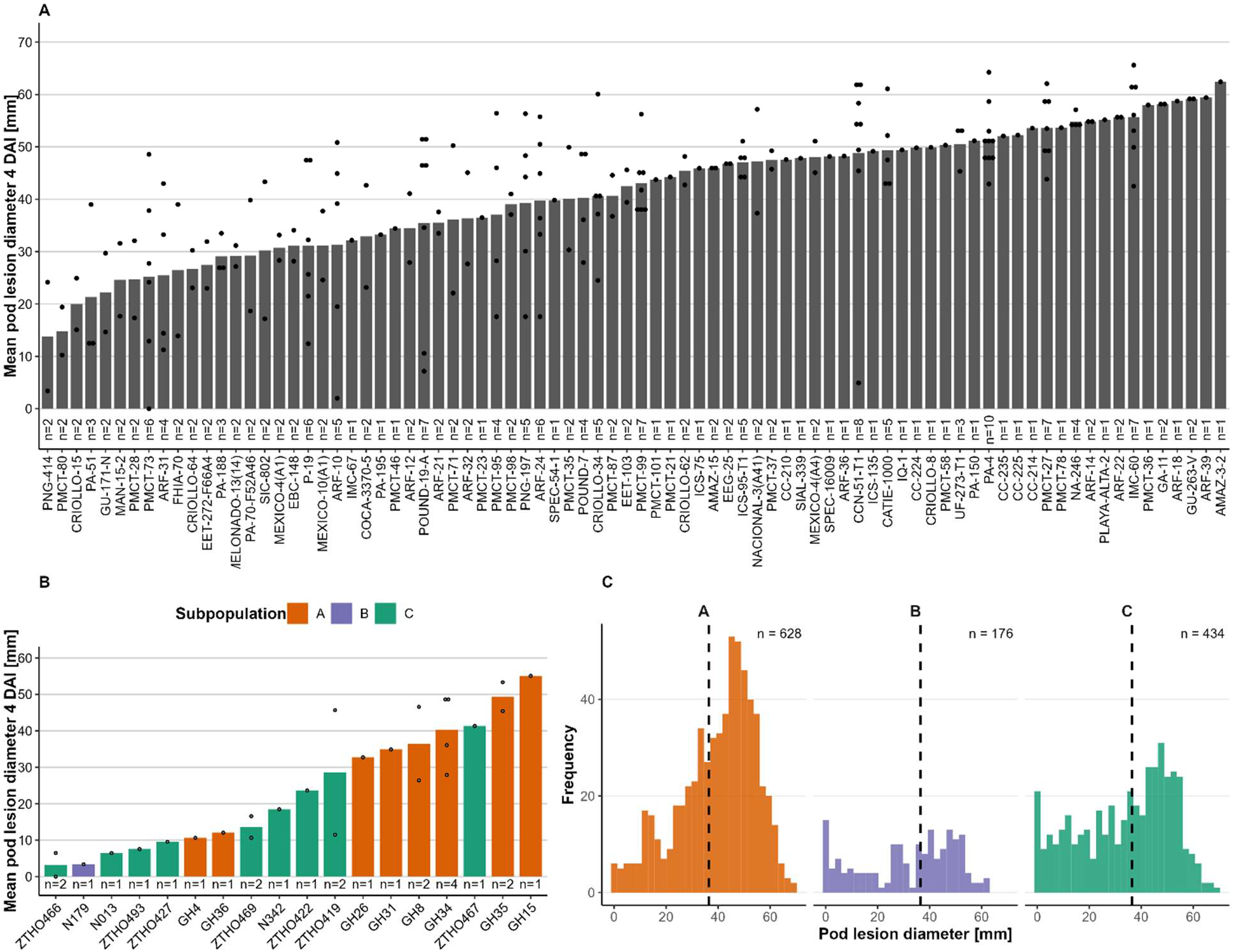
Overview of the disease screening carried out in this study. **(A)** Disease screening using the *P. megakarya* isolate GH34 on different cacao clones. **(B)** Disease screening of the cacao clone POUND-7 inoculated with different *P. megakarya* isolates. The bar plots are colored by subpopulation. **(C)** Histograms summarizing the phenotypic data, with the mean pod lesion diameter per isolate separated by subpopulation.

### Host genomic relationships improve prediction of lesion size

A series of six increasingly complex mixed models was compared to evaluate the impact of genomic information on predictions of lesion size (Table 1). These models progressively incorporated GRMs for cacao clones (Model 2), *P. megakarya* isolates (Model 3), both together (Model 4), and a dual-genome similarity matrix used to model clone-by-isolate interactions (Model 5). Model 1 attributed 0.20 of the phenotypic variance to differences among clones and 0.36 to differences among isolates, whereas clone-by-isolate interaction accounted for 0.02. Both type-B correlations were high. Model 2 replaced the traditional clone matrix with GRM_c_. In Model 2, the cacao genomic component accounted for 27% of phenotypic variance and model fit improved relative to Model 1, highlighting the value of accounting for genetic relatedness among cacao clones. Model 3 incorporated isolate relatedness while using an identity matrix for clones. This led to a slightly increased pathogen variance estimate (0.38) compared to Model 1 and a similar clone heritability estimate (0.20). Together with a slightly worse model fit, this suggests that isolate-relatedness contributed little to explaining lesion size variation. Model 4 combined genetic relatedness for both clones and isolates. This model improved fit compared to Model 1, but not compared to Model 2, with an increased clone heritability (0.26), a similar isolate variance estimate (0.36), and a smaller error term (0.38). The clone-by-isolate interaction term is at the boundary in Model 4. Model 5 added the dual-genome similarity matrix to explicitly model genomic similarities in clone-by-isolate interactions. This model yielded values similar to those of Model 4 across all parameters and a clone-by-isolate estimate of 0.01. Apart from providing a clone-by-isolate estimate, Model 5 performed like Model 4 in fit. Across these five models, Model 2 showed the best model fit with an AIC of 7503.73. Likelihood ratio tests comparing Model 2 with reduced models showed significant variance among bioassays (LR = 165.1, P < 0.001), clones (LR = 204.3, P < 0.001) and isolates (LR = 212.4, P < 0.001), whereas the clone × isolate variance was not significant (LR = 2.23, P = 0.07). All tests had one degree of freedom, with P-values from a 50:50 mixture of χ²₀ and χ²₁ (Self & Liang, 1987). Type-B correlations remained high across all models, but both are bounded functions of the interaction variance and approach unity as that variance approaches zero, so they are reported here as descriptions of the fitted variance partition rather than as evidence of stability. In clone-wise cross-validation, Model 2 predicted phenotypically masked clones within the IC3 population with a predictive ability of 0.414 ± 0.008 across five random 10-fold partitions, corresponding to an estimated prediction accuracy of 0.52. Replacing independent isolate effects with the block-diagonal pathogen GRM did not improve prediction; predictive ability differed by only 0.002 ± 0.001 in favor of Model 2.

Subpopulation means differed (Wald χ² = 9.75, 2 df, P = 0.008) with predicted lesion diameters of 40.0 ± 2.3 mm for subpopulation A, 32.6 ± 2.6 mm for B, and 34.1 ± 2.8 mm for C, and residual variances of 85.3, 107.4, and 134.5 mm² for A, B, and C.

Model 2 produced genomic estimated breeding values (GEBVs) for cacao clones and BLUPs for independent isolate effects; these are reported in Supplementary Tables 5 and 6, respectively. Clone GEBVs were centered at zero, with negative values indicating genetically smaller lesions and positive values indicating genetically larger lesions relative to the fitted population mean (35.6 mm). Predicted lesion diameter on the observed scale was obtained by combining the clone GEBV with the relevant fixed-effect mean. The higher the GEBV, the greater the potential to be susceptible; the lower the GEBV, the greater the potential to be resistant. The three isolates with the largest predicted isolate effects were PH029 (+21.6 mm), N77 (+11.5 mm), and GH12 (+10.9 mm). The three least susceptible cacao clones to *P. megakarya* infection, as phenotyped, were HY-2714182 (GEBV -18.5), COCA-3370-5 (GEBV -20.4), and LCTEEN-37 (GEBV -20.9). The three least susceptible cacao clones that were not phenotyped but were included in the cacao GRM were POUND-16-B (GEBV -17.6), NA-232 (GEBV -18.4), and NA-670 (GEBV -19.4).

Model 6 must be viewed separately from the previously described Models 1 to 5, as it introduced subpopulation-specific genomic variance components by fitting a heterogeneous correlation structure for clones across subpopulations of the *P. megakarya* isolates, along with subpopulation-specific variances for isolates. This allowed an independent estimation of variance components within each of the three *P. megakarya* subpopulations and finer biological resolution. The heterogeneous correlation structure was of particular interest as it provides estimates of the genomic correlations between subpopulations and indicates the stability of resistance across cacao clones. The estimate of 0.86 indicates strong concordance in cacao resistance predictions across subpopulations of *P. megakarya*, suggesting that population-specific resistance mechanisms were not observed. The proportion of phenotypic variance attributable to cacao genomic effects ranged from 0.19 in subpopulation B to 0.30 in subpopulation A, and 0.33 in subpopulation C. The corresponding isolate variance ratios were 0.40 in A, 0.49 in B, and 0.26 in C. These estimates indicate heterogeneity in both host genomic and isolate variance across pathogen subpopulations, although standard errors of the estimates are large. Model 6 had the lowest AIC of all models (7491.3, against 7503.7 for Model 2), indicating an improvement in fit. Model 6 provided comparisons of genetic control and resistance predictions for each subpopulation. Because isolate GH34 was sampled in Ghana and subpopulation A was larger, 51% of the phenotype data were collected from this subpopulation. This compares with the subpopulations B and C, which accounted for 33% and 15% of the phenotypic data, respectively (Figure 4C).

The three most resistant clones for each *P. megakarya* subpopulation and for Model 5 showed some overlap (Supplementary Table 5). In each GEBV list, LCTEEN-37 and COCA-3370-5 were among the three most resistant phenotyped clones, and genomic predictions identified NA-670 and NA-232 as the three most resistant non-phenotyped clones. Model 5 and GEBV of subpopulation B of Model 6 also shared HY-2714182 as a top 3 phenotyped clone, and GEBV of subpopulations B and C from Model 6 shared NA-710 as a top 3 non-phenotyped clone. The additional top 3 clones with predicted resistance were IMC-27 (GEBV subpopulation C) and PNG-414 (GEBV subpopulation A), both with phenotype information, and NA-807 (GEBV subpopulation A), without phenotype information.

To explore whether predicted resistance covaried with cacao ancestry, Model 2 GEBVs of the 350 phenotyped clones were regressed jointly on their STRUCTURE ancestry proportions. Ancestry proportions accounted for 26% of the variance in GEBVs among the 350 phenotyped clones (adjusted R² = 0.26). Relative to ‘Amelonado’, ancestry from ‘Nanay’ (−13.4 ± 1.9 mm per unit ancestry, *p* < 0.001), ‘Contamana’ (−11.0 ± 2.4, *p* < 0.001), ‘Purus’ (−8.7 ± 2.6, *p* = 0.001) and ‘Maranon’ (−4.1 ± 1.4, *p* = 0.003) was associated with smaller predicted lesions. Ancestry from ‘Criollo’ (+6.2 ± 2.4, *p* = 0.010) and ‘Guiana’ (+4.1 ± 1.7, *p* = 0.013) was associated with larger lesions (Figure 5). Using this compositional regression, replacing one-half unit of ‘Amelonado’ ancestry with ‘Nanay’ ancestry corresponds to an expected GEBV decrease of approximately 6.7 mm. Coefficients for ‘Iquitos’ and ‘Nacional’ ancestry were not distinguishable from zero in this exploratory regression, whereas ‘Curaray’ and ‘Bolivian’ ancestry were too rare among phenotyped clones for reliable estimation. Variance inflation factors were moderate, ranging from 1.0 to 1.5.

**Figure 5:**
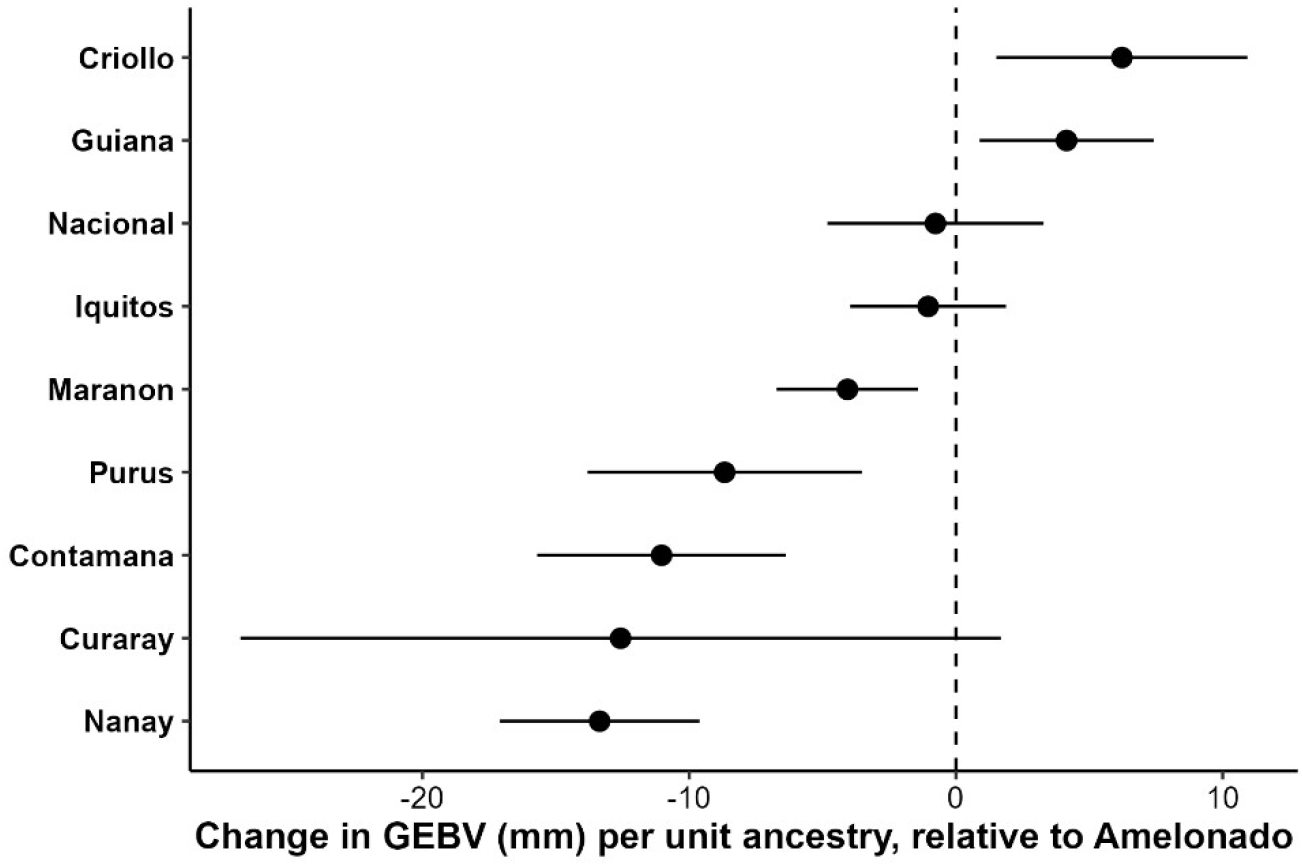
Association between genetic-group ancestry and the genomic estimated breeding values (GEBV) of 350 phenotyped cacao clones (Model 2). Points are regression coefficients from a multiple regression of GEBV on STRUCTURE ancestry proportions, with ‘Amelonado’ as reference; bars are 95% confidence intervals. A coefficient of −10 means that a clone of entirely that ancestry is predicted to have lesions 10 mm smaller than an ‘Amelonado’ clone. Negative values indicate smaller predicted lesions, that is, higher resistance. ‘Bolivian’ is omitted because too few phenotyped clones carry this ancestry.

## Discussion

### A sparse screen as a first map of resistance to the emerging pathogen *P. megakarya*

When this screen began, quantitative differences in aggressiveness among diverse *P. megakarya* isolates, and the extent to which such variation affected genomic prediction of host resistance were poorly characterized. We therefore sampled broadly across the genetically diverse IC3 collection rather than restricting screening to a small set of previously characterized resistance sources. To our knowledge, this is the largest detached-pod screen against *P. megakarya* in number of isolates (58) and one of the largest in number of clones (350 genotyped clones). This screening provides the first genomic predictions of resistance to this pathogen for the entire IC3 germplasm collection at CATIE. Three findings define the principal results of this study. First, genomic relationships among cacao clones improved model fit and enabled prediction of resistance for all clones in the IC3 collection. Second, *P. megakarya* isolates differed substantially in aggressiveness, and broad pathogen subpopulations differed in mean lesion severity, while the inclusion of genomic relationships among isolates did not improve predictions of resistance. Third, clone rankings were largely concordant across pathogen subpopulations.

### Host relatedness improves fit and prediction

Ranking the models by AIC gave a consistent picture (Table 1). Subpopulation-specific variances provided the model that fit the data best (Model 6), followed by inclusion of the cacao relationship matrix alone (Model 2). Adding the pathogen relationship matrix, alone (Model 3) or as an interaction term incorporating both relationship matrices (Models 4 and 5), did not improve fit. Incorporating the cacao GRM increased the proportion of phenotypic variance attributed to the host genomic component from 0.20 to 0.27 and improved model fitting, consistent with genomic relationships recovering information that was otherwise unavailable for sparsely phenotyped clones. Clone-wise cross-validation showed that this information can be transferred to phenotypically masked clones within the related IC3 population (predictive ability 0.41; estimated accuracy 0.52), a level comparable to host predictions in the maize–*Fusarium verticillioides* dual-genome study (Hudson et al., 2024). Romero Navarro et al. (2017) reported accuracies of about 0.5 for black pod in cacao using means of eight years of field observations under natural *P. palmivora* infection. Because clones were assigned to folds at random, validation clones generally retained genomically related individuals in the training set. The resulting accuracy therefore describes interpolation within IC3, and further validation is required before extrapolating model predictions into other populations.

The analysis illustrates the utility of relationship-matrix scaling when comparing variance components across different populations. Before the relationship matrices were rescaled, models with the pathogen matrix attributed 59–61% of phenotypic variance to isolates, nearly twice the estimate from models treating isolates as unrelated. This inflation resulted from differences in GRM scale rather than evidence for substantially greater pathogen genetic variance. In this predominantly clonal diploid pathogen, a large proportion of retained loci were fixed heterozygous within subpopulations, resulting in GRMs whose diagonal scale differed strongly from that of cacao. Scaling each matrix to a common base (Legarra, 2016) removed the discrepancy and suggests that pathogen relationship matrices be reported along with their scale.

### Isolates differ, but within-subpopulation genomic relatedness does not explain their aggressiveness

Differences among isolates explained more lesion-size variation than differences among clones (0.34 against 0.27 of phenotypic variance in Model 2), and in this dataset subpopulation A produced lesions 6–7 mm larger than subpopulations B and C. Working with eight isolates, Ali et al. (2016) found *P. megakarya* uniformly aggressive on pod husks; our larger panel shows that isolates of this species do differ substantially. After accounting for mean differences among pathogen subpopulations, incorporating within-subpopulation genomic relationships among isolates did not improve model fit or prediction accuracy. The substantial variation among isolate effects was not well represented by genome-wide relatedness at the captured loci.

At least three features of the dataset and pathosystem may explain this result. First, *P. megakarya* reproduces mainly clonally (Nyassé et al., 1999), and 84–92% of captured SNPs were fixed heterozygous within subpopulations, leaving 882–5,309 informative SNPs within the virulence-associated genes selected for sequencing. Genome-wide relatedness at the retained loci may primarily reflect clonal descent and broad genomic similarity, rather than variation in gene content or expression differences that underlie aggressiveness. Second, about 84% of *P. megakarya* genes are duplicated after a recent whole-genome duplication (Morales-Cruz et al., 2020). Collapsed or highly similar paralogous sequences can generate apparent heterozygosity when reads from different gene copies map to a single reference locus. Consequently, some of the observed fixed heterozygosity may reflect sequence differences between paralogs rather than allelic variation. Although genuine genetic variation among individuals may still be detectable at these loci, collapsing paralogous sequences can mask allelic differences or distort genotype calls, potentially leading to an underestimation of genetic diversity. Third, isolate effects were estimated with error, because 51% of isolate pairs could not be compared on the same clone and the control isolate GH34 alone was used to inoculate 227 pods. Part of the isolate variance may also be non-genetic, reflecting culture history or inoculum quality, so parameter estimates for the pathogen are reported as the share of phenotypic variance rather than as a genetic parameter estimate for the ‘heritability’ of pathogen virulence.

Previous dual-genome studies likewise found limited improvement in host prediction after incorporating pathogen genomic relationships, although the pathogen biology and population structures differed substantially from those of *P. megakarya* (Domínguez-Rondón et al., 2025; Hudson et al., 2024). Other pathogen populations studied were haploid ascomycetes with weak population structure and regular (*Zymoseptoria tritici*) (Morais et al., 2019) or occasional (*F. verticillioides*) (Fallahi et al., 2021) recombination. Our study therefore tested the approach using a diploid, clonally reproducing oomycete with strong geographic structure.

The differences between subpopulations need one caveat. Our isolates were assembled from collections made for different purposes, and isolates sampled at an epidemic front may be more aggressive than those baited from soil or other parts of the tree; without consistent sample metadata, intrinsic aggressiveness cannot be separated from sampling origin. The broad west-to-east genomic structure observed here is consistent with previously described geographic differentiation of *P. megakarya* populations across West and Central Africa by Nyassé et al. (1999), including intermediate genotypes near the Nigeria–Cameroon border. The larger number of segregating SNPs observed in subpopulation C is consistent with greater diversity in populations in Cameroon, although mapping all isolates to the Ghanaian GH34 reference could also inflate variation in more divergent lineages. The sequence data may provide a resource for further studies of the diversity of virulence associated in *P. megakarya*. Subpopulations differed in which effector and protease genes carried SNPs (Figure 3), and a paralogue-aware analysis of effector presence–absence variation may be used to identify isolates with different virulence gene repertoires. Screening cacao populations with a few diverse isolates would improve the odds of identifying isolate or effector-specific resistance mechanisms within cacao collections.

### Concordant clone rankings, unresolved interaction

The genomic correlation of clone effects across pathogen subpopulations (0.86 ± 0.06) indicates substantial concordance in host rankings across the three modeled pathogen groups. Unlike type-B correlations, which approach unity whenever the interaction variance is small, this correlation is estimated as a parameter in its own right. This agrees with earlier evidence of broadly consistent resistance rankings across *P. palmivora* isolates (Surujdeo-Maharaj et al., 2001), between *P. palmivora* and *P. megakarya* (Nyadanu et al., 2012) and across countries (Nyassé et al., 2007). Exceptions exist: SCA 6 restricts *P. palmivora* but is invaded by *P. megakarya* (Ali et al., 2016; Herrera-Corzo et al., 2026), so resistance to one species does not guarantee resistance to the other.

The fitted clone-by-isolate interaction accounted for only 0.02 of phenotypic variance and was not statistically distinguishable from zero (P = 0.07). Given the sparse connectivity of the design, however, this result should be interpreted as limited evidence for interaction rather than evidence that clone-by-isolate specificity is absent. This reflects the experimental design as much as the biology. Only 5% of possible combinations were tested, most of them once, and lesion sizes on POUND-7, tested with 18 isolates, varied in ways that a single variance component averages away. While a fully factorial screen is not required to provide estimates of interaction variance, traditional designs using a selection of diverse cacao clones and isolates would provide more precise estimates of variance components. Circular allocations of isolates to hosts, as used by Hudson et al. (2024) may also be used to increase the efficiency per observation (Huber et al., 1992) and improvements in the accuracy of predictions from sparse designs rise with the overlap of genotypes across test conditions (Atanda et al., 2022). Combining the sparse testing strategy of this study, which provides phenotype data for many clones and genomic predictions for all clones, with more traditional screening of a small set of varieties would provide both validation and evaluation of the trade-off between testing large numbers of clones and obtaining precise genetic parameter estimates.

### What lesion size on detached pods measures

The phenotype measured here represents resistance of unwounded pods (harvested at approximately four-month since formation) to penetration and early lesion expansion four days after inoculation. It captured penetration and early lesion expansion on the pod, but escape through flowering time, fruiting-cycle length, or pod position also contribute to field resistance (Efombagn et al., 2004). Pods were inoculated 4–13 days after harvest due to shipping constrants, so variation associated with post-harvest interval likely contributed to residual variation and reduced precision of clone and isolate effects. Laboratory and field rankings agree at the level of genotype means, but not always significantly (Efombagn et al., 2011), and natural field infection can be so dominated by environment that its broad-sense heritability falls to 0.15 (Nyadanu et al., 2012). Controlled assays therefore remain the practical basis for ranking genotypes, but the top-ranked clones identified here require field evaluation under *P. megakarya* pressure for definitive validation.

### Genetic groups and candidate parents

In the exploratory ancestry regression, higher proportions of ‘Nanay’, ‘Contamana’, ‘Purus’, and ‘Maranon’ ancestry were associated with smaller predicted lesion sizes. The negative ‘Maranon’ coefficient is consistent with previous reports of black-pod resistance in PA material and with lower natural black-pod incidence among clones carrying ‘Maranon’ ancestry (Iwaro et al., 2005; Romero Navarro et al., 2017). Our predictions also place an established ‘Nanay‘ source, POUND-7 (GEBV −12.2 ± 1.7 mm, 26 pods), and several unphenotyped ‘NA’ clones among the most resistant. ‘Guiana’ ancestry showed the opposite trend in the present dataset, with higher ‘Guiana’ ancestry associated with larger predicted lesions. This finding contrasts with wild ‘Guiana’ genotypes described as resistant to both *P. megakarya* (Paulin et al., 2008) and *P. palmivora* (Thevenin et al., 2012). Those studies used a single isolate per pathogen species, whereas our clones were challenged with up to four different isolates, many from the more aggressive subpopulation A; the wild French Guiana populations are not the ‘GU’ clones held at CATIE; and leaf and pod resistance overlap only partly (Iwaro et al., 1997). Not every reported source of resistance ranked highly either: PA-150, a good general combiner for resistance in Ghana (Nyadanu et al., 2012), was close to the population mean (+0.3 ± 3.7 mm). Because the response variable consisted of marker-derived, shrunken GEBVs and the phenotyped clones were a non-random, predominantly admixed subset of IC3, these ancestry associations should be interpreted as hypothesis-generating trends rather than estimates of causal genetic-group effects.

### From predictions to breeding suggestions

Validation of the genomic prediction model is needed before making strong recommendations on its application to reduce the impact of black pod disease. The experimental design enabled testing of a large number of cacao varieties, leveraging genomic connectivity across bioassays to provide unbiased predictions for any genotyped cacao clone. This approach accommodates temporal variability in pod availability and is well suited for the initial characterization of germplasm collections and variable pathogen populations. The IC3 germplasm collection is highly diverse and is likely related to ex-situ germplasm populations used around the world, providing some confidence that the IC3-trained prediction models may be applicable globally. However, prediction accuracy may decline in genetically distinct germplasm collections, as model transferability depends on differences in resistance among populations, shared genetic ancestry, and consistent linkage between genetic markers and resistance-associated loci.

A practical suggestion for responding to the black pod epidemic in Africa is to ensure clones from populations demonstrating resistance, such as ‘Nanay’, ‘Contamana’, ‘Purus’, are made available for evaluation in the region. Importation of clones with predicted resistance for field evaluation in areas with high infection and for use as parents to cross with local selections is also recommended. Genotyping African germplasm collections using the IC3-trained prediction model to identify 10 to 20 resistant and susceptible clones may be used to focus validation on clones with contrasting resistance predictions. Subsequent screening with local isolates would provide the phenotype data required for model validation in independent populations. Given the stability of cacao clone rankings across pathogen populations, phenotype and genotype data from African populations screened for resistance to local isolates may be incorporated into a general resistance prediction model that the cacao community may build upon. As isolates evolve, or if repeatable changes in rankings across cacao clones are identified, isolate genotypes may be incorporated into prediction models to account for interactions not detected in this study.

Under the average fitted pathogen background and the lesion-size thresholds used for this screening assay, 215 clones had predicted lesion diameters below 30 mm, including 119 clones without direct phenotype records. These values may be used to rank clones for validation studies that would provide more definitive resistance classifications. Their practical value is in prioritizing clones from a large germplasm collection for targeted phenotyping and crossing. Releasing a new cacao variety can take 20–30 years (Kulesza et al., 2024), and *P. megakarya* is still confined to West and Central Africa. Once validated, predictions from a quarantine screen could also help breeders in regions not yet invaded to decide which clones to test first, and the framework accounting for both host and pathogen variability can be updated as additional screening and field data accumulate.

The heterogeneous correlation structure was of particular interest as it provides estimates of the genomic correlations among subpopulations and indicates the stability of resistance across cacao clones. The estimate of 0.86 indicates strong concordance in cacao resistance predictions across *P. megakarya* subpopulations, suggesting that subpopulation-specific resistance mechanisms were not identified. Screening with isolates from different populations provided similar predictions of resistance, suggesting that these results may be combined with additional screening data from African populations to prioritize clones for the development of resistance and to expand the inferences we can draw using genomic prediction models.

## Conclusions

We combined genotyping of the CATIE International Cacao Collection with capture sequencing of *P. megakarya* and a sparse detached-pod screen to produce the first genomic predictions of resistance to black pod disease. Genomic relationships among cacao clones improved model fit and enabled prediction of phenotypically untested clones, whereas modeling within-subpopulation genomic relationships among *P. megakarya* isolates provided no measurable improvement beyond adjusting for fixed pathogen subpopulation effects. Clone rankings were broadly concordant across the three pathogen subpopulations, and exploratory ancestry analyses identified several Upper Amazon ancestry components associated with lower predicted lesion size. Replicated screening of a well-connected subset of clones and isolates, field validation of the top-ranked clones, and examination of drivers of variation in pathogen virulence are the next steps. More broadly, the study illustrates how genomic relationships can be used to connect sparse disease-screening datasets in perennial crops and how pathogen genomic covariance structures can be incorporated into resistance and virulence prediction models. Host genomic relationships can extend information from a sparse screening population to related unphenotyped germplasm, while explicit evaluation and appropriate scaling of pathogen relationship matrices are essential before attributing phenotypic variation to pathogen genomic effects.

## Acknowledgements

We would like to thank the field workers at CATIE for harvesting and providing cacao pods, and for technical assistance in producing and shipping pods; Robert Henfling for assistance in shipping pods; Dr. Brian Bailey and Dr. G. Martijn ten Hoopen for the provision of the *Phytophthora megakarya* isolates; the University of Florida for facilities and infrastructure; and Mars, Incorporated for funding this project.

## Disclosures, declarations, and project funding

Authors of this paper (DL, AF, DD, DH, DR, JPM) are or were employed by Mars, Incorporated, and the company has provided funding to CATIE and the University of Florida to complete this project.

## Supplementary Materials

**Supplementary Table 1: Geographic origin and information on the 67 Phytophthora megakarya isolates used in this study.** Low virulent = more than 80% of clones showed no infection with these isolates.

Supplementary Table 2: Summary of the gene families, the number of candidate genes, and the number of targeted genes for the capture sequencing approach used for *Phytophthora megakarya* isolates in this study. The gene annotations were obtained from Morales-Cruz et al. (2020, File_S1).

**Supplementary Table 3: Results of the STRUCTURE analysis on the 947 clones investigated in this study.**

**Supplementary Table 4: Summary of the number and gene family of the genes with genetic variation (SNPs) within each *Phytophthora megakarya* subpopulation.**

**Supplementary Table 5: Genomic estimated breeding values (GEBVs) for cacao clone resistance obtained from Model 2 and 6**. Given are the number of inoculated pods in the bioassay (inoc), the predicted breeding value for clone c (Best Linear Unbiased Prediction, BLUPc), the standard error of prediction for the breeding value of clone c (SEPc), and the standardized Wald-type test statistic Zc (calculated as BLUPc/SEPc) for Model 2. The same metrics are given for Model 6, separated in GEBVs from subpopulation A, B, and C.

**Supplementary Table 6: Genomic estimated breeding values for isolate virulence obtained from Model 2.** Given are the predicted breeding value for isolate i (Best Linear Unbiased Prediction, BLUPi), the standard error of prediction for the breeding value of isolate i (SEPi), the standardized Wald-type test statistic Zi (calculated as BLUPi/SEPi), the number of inoculated pods in the bioassay (inoc), and the assignment of a subpopulation (subpop), either A, B, or C.

**Supplementary Table 7: Raw variance components (mm² ± SE) of each model.**

**Supplementary Figure 1: Population structure of 67 Phytophthora megakarya isolates. (A)** Principal component analysis (PCA) plot showing PC1 and PC2 from 62,203 SNPs across 67 *P. megakarya* isolates. Each point represents an individual isolate, colored by their country of origin.**(B)** Correlation matrix of 67 P. megakarya isolates based on marker data. The color bar to the left of the plot shows the country of origin of each isolate.

**Supplementary Figure 2. Additional principal component projections of the genetic structure of 947 clones of the IC3 cacao collection.** Principal component analysis (PCA) was based on 8,572 SNP markers. **(A)** Projection of principal components 2 and 3, explaining 12.0% and 7.6% of the genetic variation, respectively. **(B)** Projection of principal components 3 and 4, explaining 7.6% and 5.7% of the genetic variation, respectively. Each point represents an individual clone. Genetic-group assignment was based on the ancestry component with the highest STRUCTURE membership coefficient. Clones with ≥75% membership in their predominant genetic group were classified as members and are shown in color, whereas clones with <75% membership were classified as admixed and are shown in gray. Genetic-group nomenclature follows Motamayor et al. (2008).

**Supplementary Figure 3: Population structure of 41 Phytophthora megakarya isolates from Ghana and Nigeria. (A)** Principal component analysis (PCA) plot showing PC1 and PC2 from 12,342 SNPs across 41 *P. megakarya* isolates. Each point represents an individual isolate, colored by their country of origin. **(B)** Correlation matrix of 41 *P. megakarya* isolates based on marker data. The color bar to the left of the plot shows the country of origin of each isolate.

**Supplementary Figure 4: (A) Population structure of 26 *Phytophthora megakarya* isolates from Cameroon and Nigeria. (A)** Principal component analysis (PCA) plot showing PC1 and PC2 from 44,633 SNPs across 26 *P. megakarya* isolates. Each point represents an individual isolate colored by their country of origin. **(B)** Correlation matrix for 26 *P. megakarya* isolates based on marker data. The color bar to the left of the plot shows the country of origin of each isolate.

Supplementary Figure 5: Population structure of the three final three *P. megakarya* subpopulations defined in this study. (A, C, E) Principal component analysis (PCA) plots showing PC1 and PC2 from subpopulation A (882 SNPs across 23 isolates), subpopulation B (995 SNPs across 14 isolates), and subpopulation C (5,309 SNPs across 21 isolates). Each point represents an individual isolate colored by their country of origin. **(B, D, F)** Heatmap of the genetic relationship matrices (GRM) of the subpopulation A (B), B (D), and C (F).

**Supplementary Figure 6: Virulence of P. megakarya isolates from three genetically distinct subpopulations across cacao clones, based on disease screening data.** Summary of all lesion size assessments for each isolate across clones. Each bar plot shows the average pod-lesion diameter across all measurements for an isolate. Standard deviation is depicted through a black line if several observations were made. Each *P. megakarya* isolate was used to inoculate a separate set of cacao clones. The isolate GH34 was used as a control (n = 209).

Supplementary Figure 7: Disease screening of cacao pods from clones of the IC3 cacao collection inoculated with different *P. megakarya* isolates. The barplots show the overall mean pod lesion diameter for each clone across P. megakarya isolates **(A)** and by isolates from subpopulations A **(B)**, B **(C)**, and C **(D)**. Standard deviation is depicted through a black line if several observations were made. The horizontal dashed line at 15 mm and 30 mm represent the thresholds for calling a clone resistant (0-15 mm), moderately resistant (15-30 mm), or susceptible (>30 mm).

