## Supplementary Figures and Tables for "Genomic prediction of resistance to *Phytophthora megakarya* in cacao from a sparse host-pathogen screen"

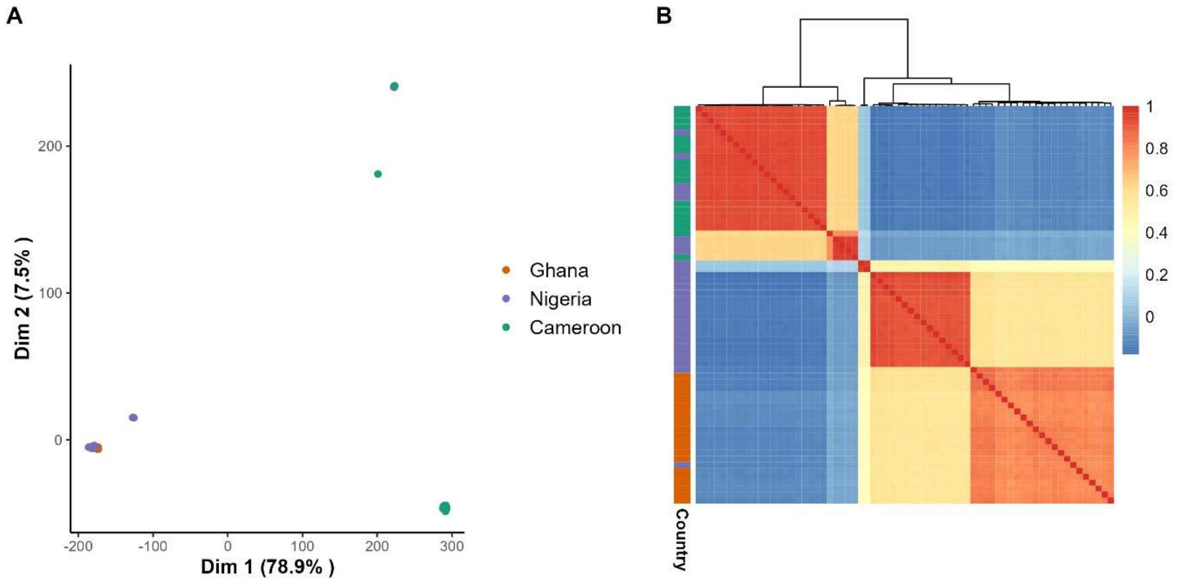


**Supplementary Figure 1: Population structure of 67 *Phytophthora megakarya* isolates. (A**) Principal component analysis (PCA) plot showing PC1 and PC2 from 62,203 SNPs across 67 *P. megakarya* isolates. Each point represents an individual isolate, colored by their country of origin. **(B)** Correlation matrix of 67 P. megakarya isolates based on marker data. The color bar to the left of the plot shows the country of origin of each isolate.

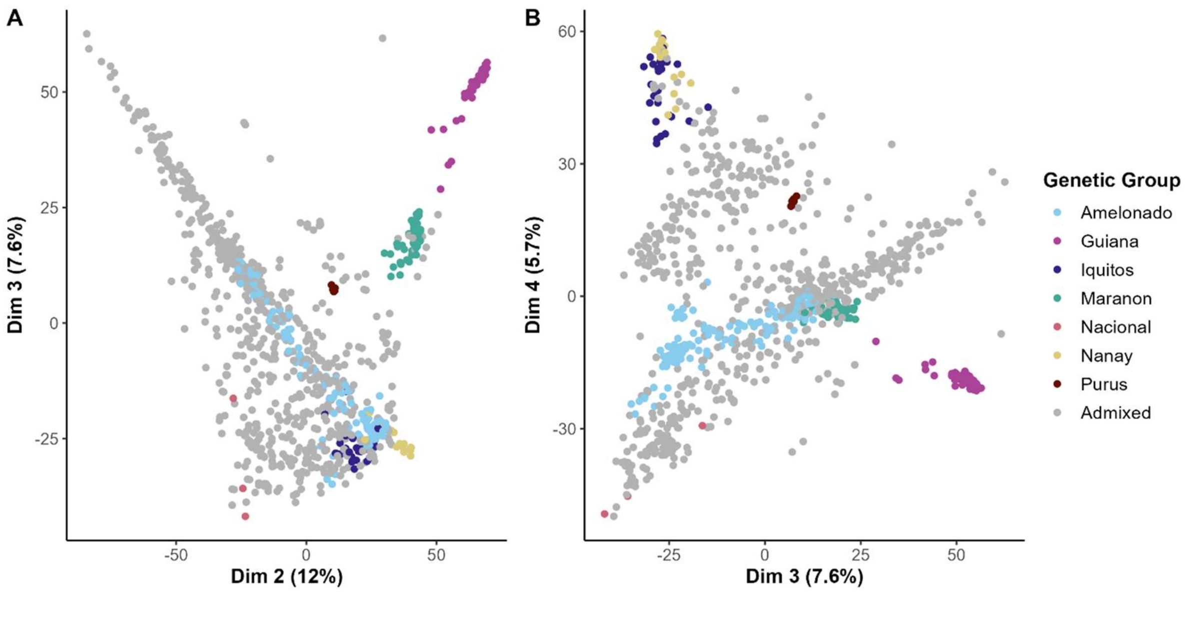


**Supplementary Figure 2. Additional principal component projections of the genetic structure of 947 clones of the IC3 cacao collection.** Principal component analysis (PCA) was based on 8,572 SNP markers. **(A)** Projection of principal components 2 and 3, explaining 12.0% and 7.6% of the genetic variation, respectively. **(B)** Projection of principal components 3 and 4, explaining 7.6% and 5.7% of the genetic variation, respectively. Each point represents an individual clone. Genetic-group assignment was based on the ancestry component with the highest STRUCTURE membership coefficient. Clones with ≥75% membership in their predominant genetic group were classified as members and are shown in color, whereas clones with <75% membership were classified as admixed and are shown in gray. Genetic-group nomenclature follows Motamayor et al. (2008).

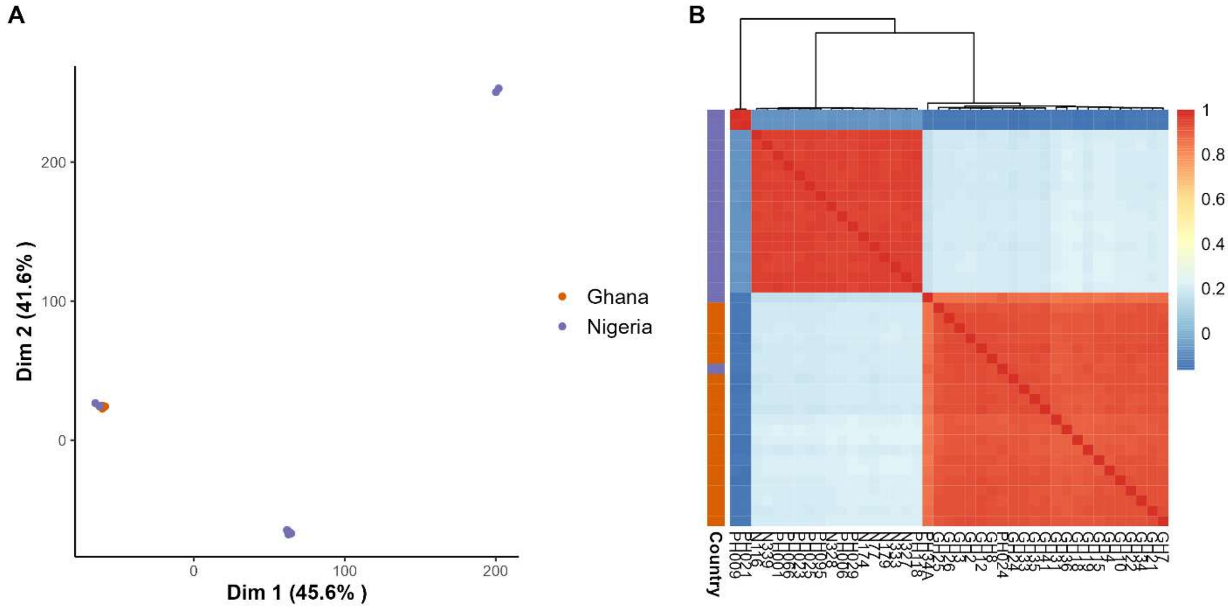


**Supplementary Figure 2: Population structure of 41 *Phytophthora megakarya* isolates from Ghana and Nigeria. (A**) Principal component analysis (PCA) plot showing PC1 and PC2 from 12,342 SNPs across 41 *P. megakarya* isolates. Each point represents an individual isolate, colored by their country of origin. **(B)** Correlation matrix of 41 *P. megakarya* isolates based on marker data. The color bar to the left of the plot shows the country of origin of each isolate.

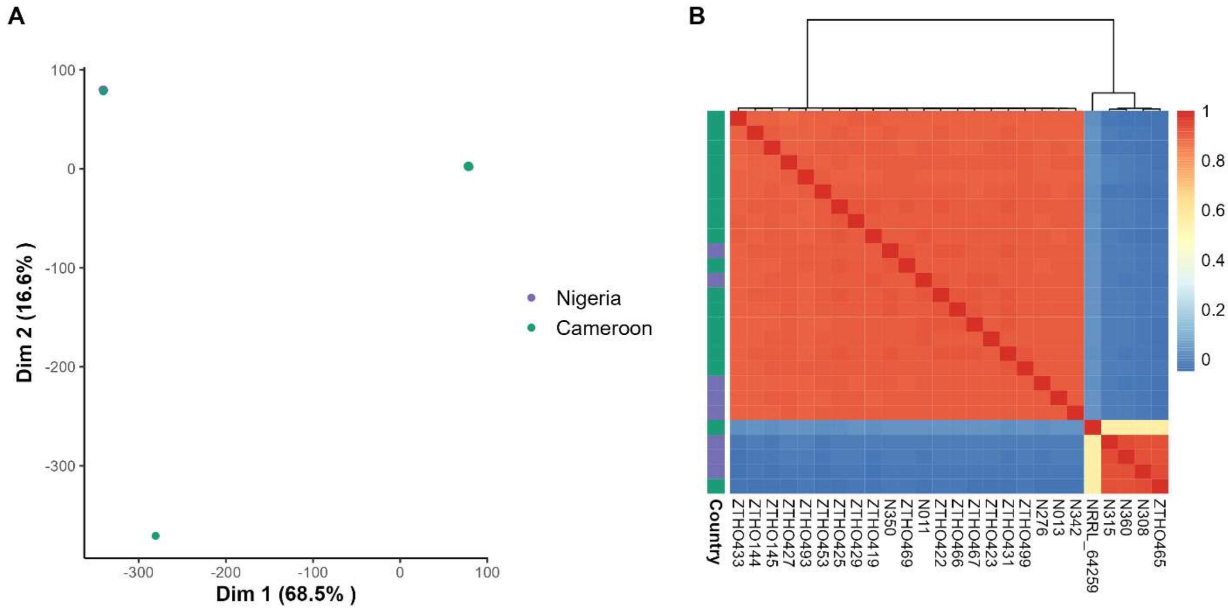


**Supplementary Figure 3: (A) Population structure of 26 *Phytophthora megakarya* isolates from Cameroon and Nigeria. (A**) Principal component analysis (PCA) plot showing PC1 and PC2 from 44,633 SNPs across 26 *P. megakarya* isolates. Each point represents an individual isolate colored by their country of origin. **(B)** Correlation matrix for 26 *P. megakarya* isolates based on marker data. The color bar to the left of the plot shows the country of origin of each isolate.

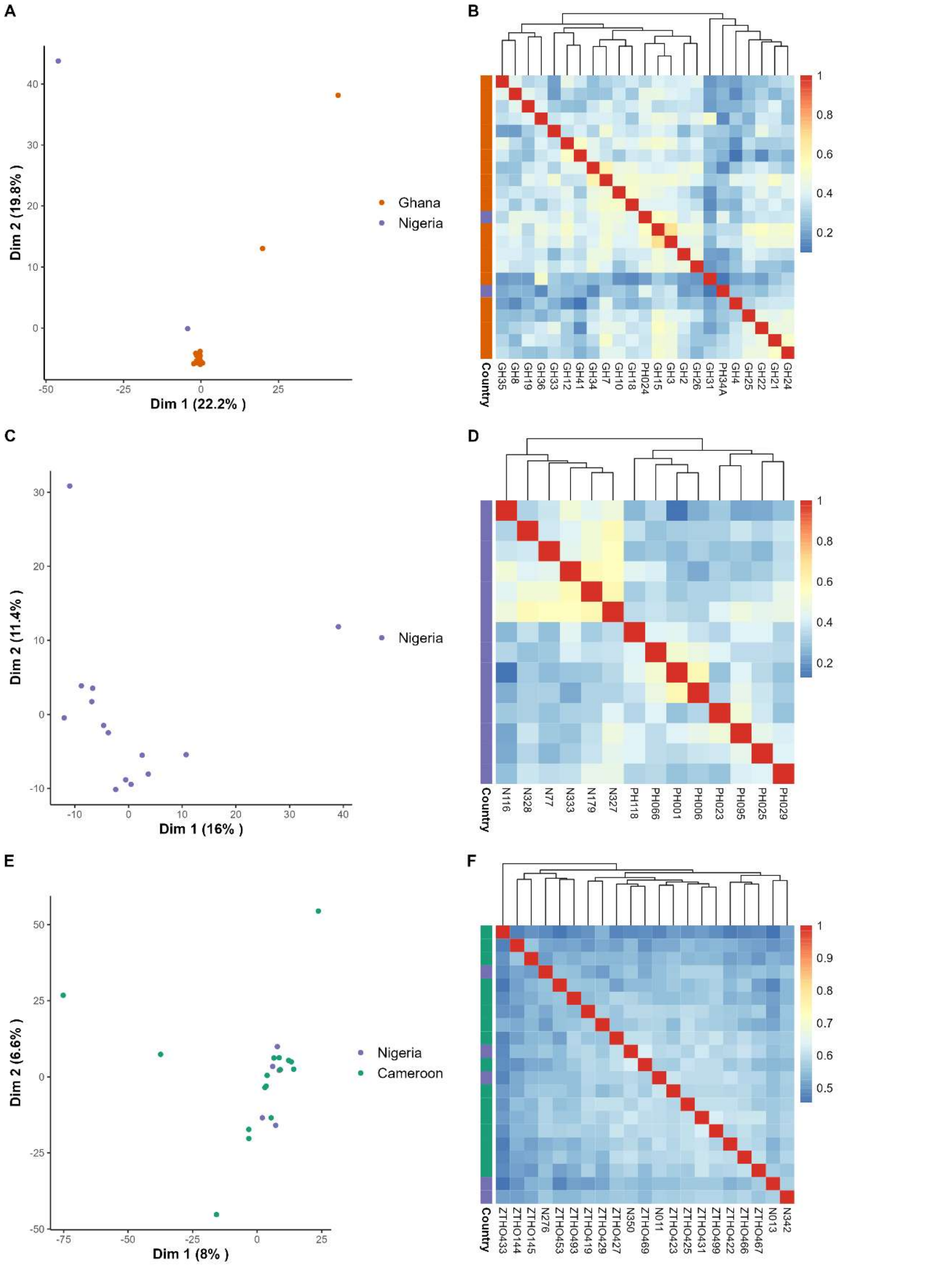


**Supplementary Figure 4: Population structure of the three final three *P. megakarya* subpopulations defined in this study. (A, C, E)** Principal component analysis (PCA) plots showing PC1 and PC2 from subpopulation A (882 SNPs across 23 isolates), subpopulation B (995 SNPs across 14 isolates), and subpopulation C (5,309 SNPs across 21 isolates). Each point represents an individual isolate colored by their country of origin. **(B, D, F)** Heatmap of the genetic relationship matrices (GRM) of the subpopulation A (B), B (D), and C (F).

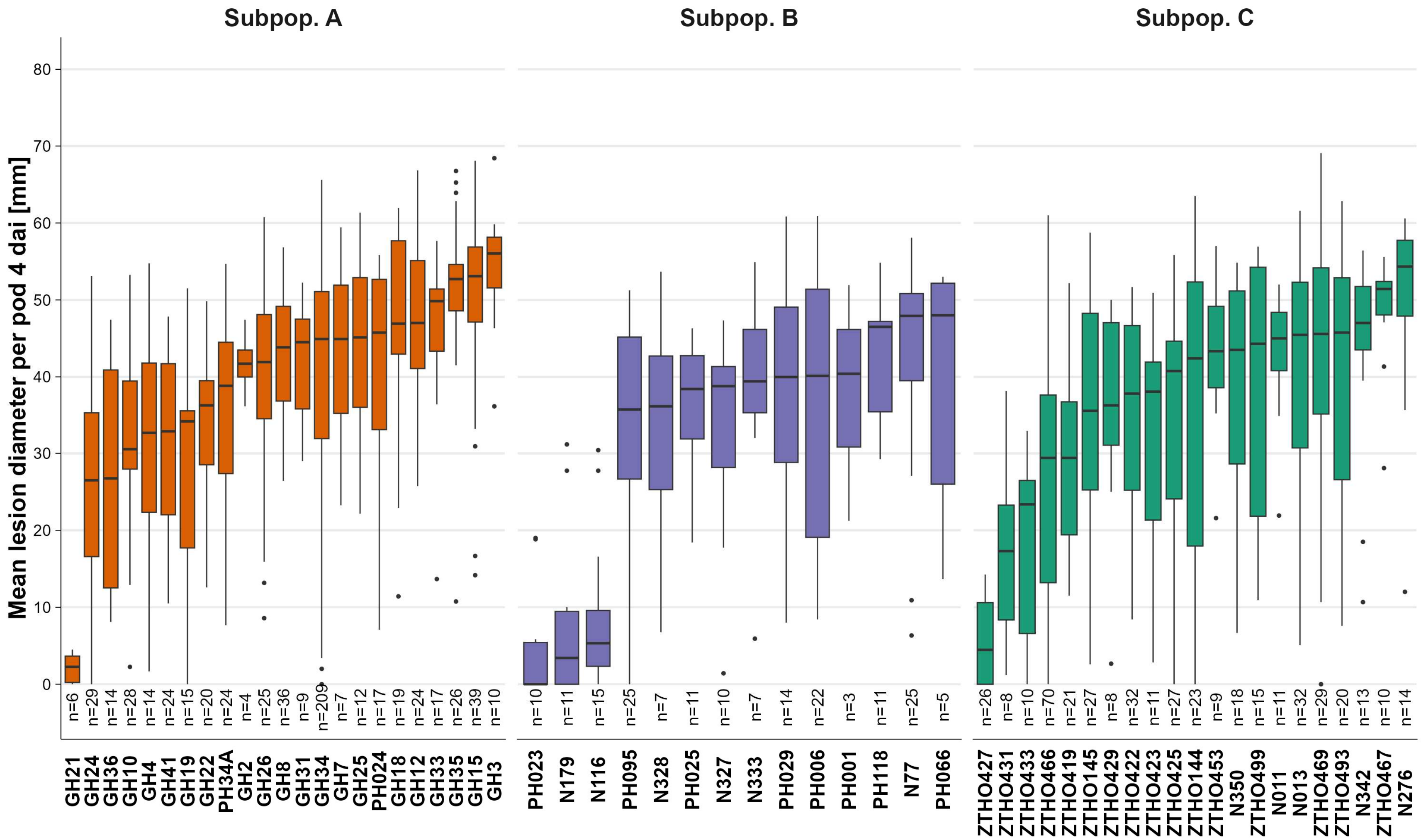


**Supplementary Figure 5: Virulence of *P. megakarya* isolates from three genetically distinct subpopulations across cacao clones, based on disease screening data.** Summary of all lesion size assessments for each isolate across clones. Each bar plot shows the average pod-lesion diameter across all measurements for an isolate. Standard deviation is depicted through a black line if several observations were made. Each *P. megakarya* isolate was used to inoculate a separate set of cacao clones. The isolate GH34 was used as a control (n = 209).

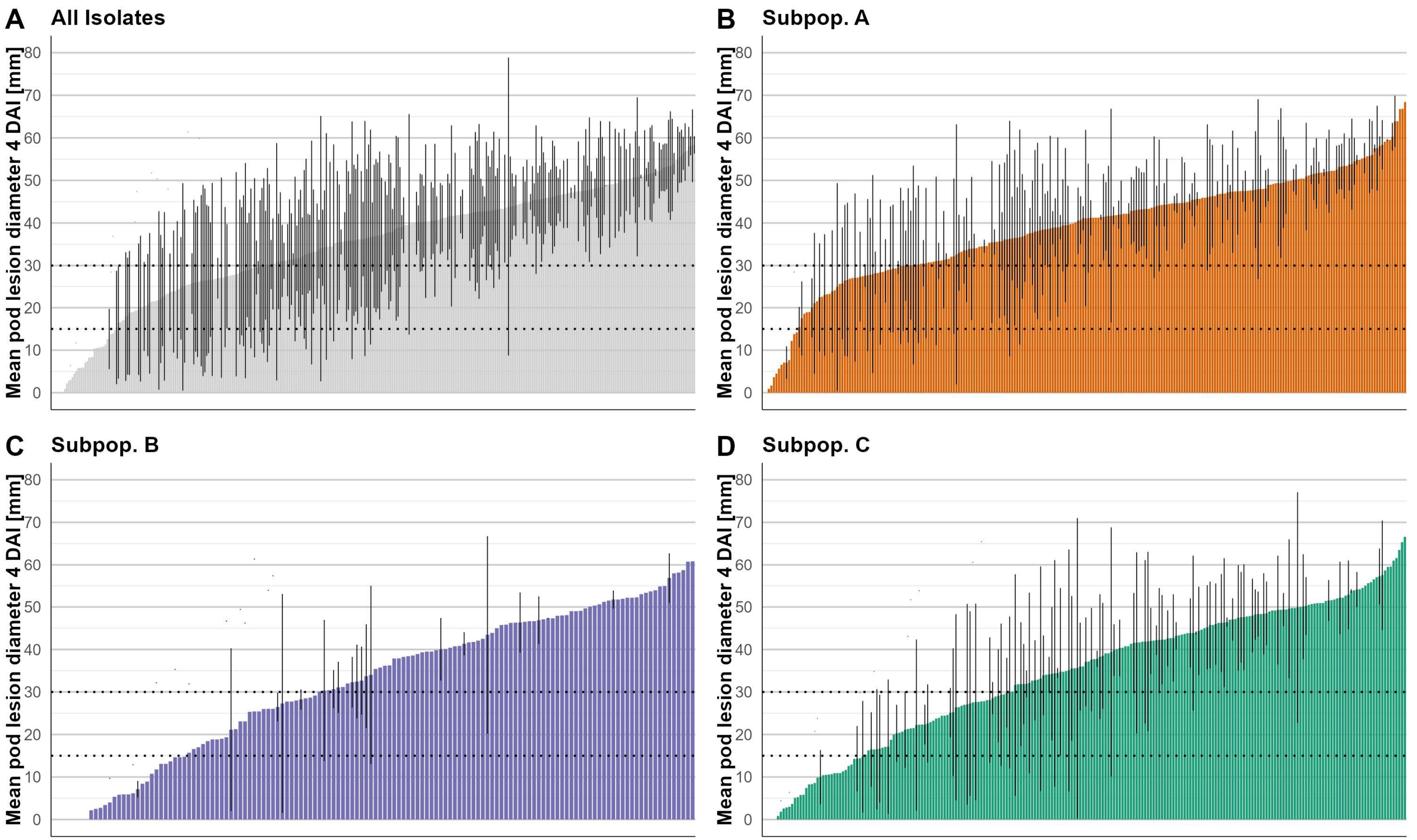


**Supplementary Figure 6: Disease screening of cacao pods from clones of the IC3 cacao collection inoculated with different *P. megakarya* isolates.** The barplots show the overall mean pod lesion diameter for each clone across *P. megakarya* isolates (A) and by isolates from subpopulations A (B), B (C), and C (D). Standard deviation is depicted through a black line if several observations were made. The horizontal dashed line at 15 mm and 30 mm represent the thresholds for calling a clone resistant (0-15 mm), moderately resistant (15-30 mm), or susceptible (>30 mm).

**Supplementary Table 7: Raw variance components (mm$^2$ ± SE) of each model.**

| Model | bioassay | clone | isolate | interaction | residual subpop. A | residual subpop. B | residual subpop. C |
|---|---|---|---|---|---|---|---|
| **1** | 48.94 ± 15.89 | 51.93 ± 7.03 | 94.37 ± 21.62 | 4.22 ± 6.67 | 84.12 ± 7.61 | 108.20 ± 16.49 | 138.09 ± 13.44 |
| **2** | 50.69 ± 16.39 | 78.10 ± 12.72 | 100.62 ± 22.72 | 4.92 ± 4.03 | 85.30 ± 6.54 | 107.35 ± 14.90 | 134.51 ± 11.78 |
| **3** | 53.62 ± 17.38 | 52.09 ± 7.00 | 101.27 ± 23.70 | 3.08 ± 6.57 | 85.03 ± 7.43 | 109.47 ± 16.66 | 139.33 ± 13.55 |
| **4** | 56.52 ± 18.22 | 77.81 ± 12.61 | 108.60 ± 25.06 | Bound | 88.92 ± 6.04 | 113.46 ± 14.39 | 141.45 ± 11.18 |
| **5** | 56.46 ± 18.21 | 77.71 ± 12.63 | 108.42 ± 25.04 | 2.31 ± 4.31 | 87.63 ± 6.54 | 111.06 ± 15.00 | 138.41 ± 12.01 |
| **6** | 56.92 ± 18.18 | **A** 83.16 ± 15.14<br>**B** 66.44 ± 22.02<br>**C** 98.34 ± 20.78 | **A** 110.40 ± 41.34<br>**B** 170.47 ± 81.65<br>**C** 78.71 ± 31.09 | NA | 85.57 ± 5.96 | 111.38 ± 15.33 | 123.15 ± 10.50 |
